# Oligodendrocyte calcium signaling sculpts myelin sheath morphology

**DOI:** 10.1101/2023.04.11.536299

**Authors:** Manasi Iyer, Husniye Kantarci, Nicholas Ambiel, Sammy W. Novak, Leonardo R. Andrade, Mable Lam, Alexandra E. Münch, Xinzhu Yu, Baljit S. Khakh, Uri Manor, J. Bradley Zuchero

## Abstract

Myelin is essential for rapid nerve signaling and is increasingly found to play important roles in learning and in diverse diseases of the CNS. Morphological parameters of myelin such as sheath length and thickness are regulated by neuronal activity and can precisely tune conduction velocity, but the mechanisms controlling sheath morphology are poorly understood. Local calcium signaling has been observed in nascent myelin sheaths and can be modulated by neuronal activity. However, the role of calcium signaling in sheath formation and remodeling is unknown. Here, we used genetic tools to attenuate oligodendrocyte calcium signaling during active myelination in the developing mouse CNS. Surprisingly, we found that genetic calcium attenuation did not grossly affect the number of myelinated axons or myelin thickness. Instead, calcium attenuation caused striking myelination defects resulting in shorter, dysmorphic sheaths. Mechanistically, calcium attenuation reduced actin filaments in oligodendrocytes, and an intact actin cytoskeleton was necessary and sufficient to achieve accurate myelin morphology. Together, our work reveals a novel cellular mechanism required for accurate CNS myelin formation and provides mechanistic insight into how oligodendrocytes may respond to neuronal activity to sculpt myelin sheaths throughout the nervous system.

## INTRODUCTION

Myelin, the spirally-wrapped, lipid-rich substance made by oligodendrocytes in the central nervous system (CNS), is critical for the rapid propagation of action potentials by neurons^1^. Myelin loss in diseases like multiple sclerosis or after injury causes severe disability^2^. In addition to its traditional role in increasing conduction velocity, the generation of new myelin and the remodeling of existing myelin sheaths has been increasingly implicated in supporting memory acquisition and learning^3–9^. Normal developmental myelin formation and its remodeling in the context of plasticity are thus essential for a properly functioning nervous system.

To generate myelin, oligodendrocyte precursor cells (OPCs) terminally differentiate and make enormous changes to their cell shape to transform into myelinating oligodendrocytes; these cells must select, ensheath, and then spirally wrap around axons while concurrently longitudinally extending myelin along many different axons. Myelin sheath morphology (e.g. sheath length, myelin thickness) is likely critical for precisely tuning conduction velocity to regulate neuronal network function. During development, sheath length and thickness are dictated, at least in part, by properties of the underlying neurons including axon diameter, neuron type, and neuronal activity (reviewed in ^10^). In adults, length and thickness of existing myelin sheaths can be dynamically adjusted by experience or experimentally-induced neuronal activity^3, 4, 9, 11, 12^, and these changes are predicted to be sufficient to significantly alter axonal conduction velocity^3, 13^. Together, myelin is likely to play a profoundly important role in regulating neural circuit activity—and the morphology of myelin sheaths is central to this function. However, how myelin sheath morphology is regulated is still an area of active study.

Myelin is composed mostly of cellular membrane. Because of this, central to myelination is the ability of oligodendrocytes to expand their membranes in precisely the right locations— around and along axons but not out into the neuropil or around non-axonal targets—and to generate sheaths with optimal lengths and thicknesses to support circuit function. To orchestrate the dramatic morphological transformations required for myelin generation, oligodendrocytes employ a myriad of cell biological processes^14^. Membrane addition by exocytosis^15–17^, cytoskeletal dynamics and cell adhesion^18–20^, and external attractive/repulsive cues^21^ all likely collaborate to properly shape myelin sheaths. How are these different cellular processes coordinated to allow myelin sheath morphology to be adjusted to neuronal properties?

Calcium (Ca^2+^) signaling is a likely candidate to regulate oligodendrocyte cell biology during myelin formation and remodeling. In other cell types, calcium signaling plays essential roles in regulating numerous cell biological processes relevant to myelination, including cytoskeletal dynamics, exocytosis, and gene expression^22–26^. Oligodendrocytes express numerous calcium-permeable channels and receptors, and calcium transients occur spontaneously and can be induced by a range of neurotransmitters in cultured oligodendrocytes (reviewed in ^27, 28^). In the developing zebrafish, local calcium transients occur in individual sheaths, and can be induced by neuronal activity^29, 30^. Intriguingly, the patterns of these calcium transients predict whether the sheaths will elongate or retract, suggesting that calcium signaling may actively control sheath morphology^29, 30^. Local calcium transients also occur in myelin sheaths in the mouse, both during developmental myelination as well as during remyelination following demyelination^31^. However, the function of oligodendrocyte calcium signaling in myelination remains unknown. Since local calcium signaling in myelin sheaths has the potential to bridge neuronal activity to cellular processes capable of remodeling individual sheaths, answering this question may also give key insights into the mechanism and importance of sheath dynamics during learning.

Here, we used a newly-developed genetic tool—the calcium pump “CalEx”^32, 33^—to determine the role of oligodendrocyte calcium signaling during myelination of the mouse CNS. We uncover a novel cellular mechanism used by oligodendrocytes to precisely sculpt myelin sheath morphology: calcium-regulated cytoskeletal assembly in nascent sheaths. This mechanism may explain how myelin sheath geometry is precisely adjusted to neuronal properties during the development and remodeling of neural circuits.

## RESULTS

### CalEx attenuates calcium signaling without affecting oligodendrocyte survival or differentiation

To test the requirement of intracellular calcium signaling in oligodendrocytes during developmental myelination, we used a transgenic mouse model—“CalEx^flox^” (short for <u>Ca</u>lcium <u>Ex</u>trusion)—that allows Cre-dependent expression of a constitutively active plasma membrane calcium pump (hPMCA2w/b) to extrude cytoplasmic calcium in Cre-expressing cells (Fig. 1A)^32, 33^. We crossed CalEx^flox/flox^ and Cnp-Cre mice^34^ to attenuate calcium signaling specifically in pre-myelinating cells of the CNS and PNS (hereafter referred to as “OL-CalEx”). OL-CalEx mice and wildtype (WT) littermates were born in normal Mendelian frequencies and survived until adulthood, but OL-CalEx mice were significantly smaller than WT littermates (Fig. S1A-B).

**Figure 1.**
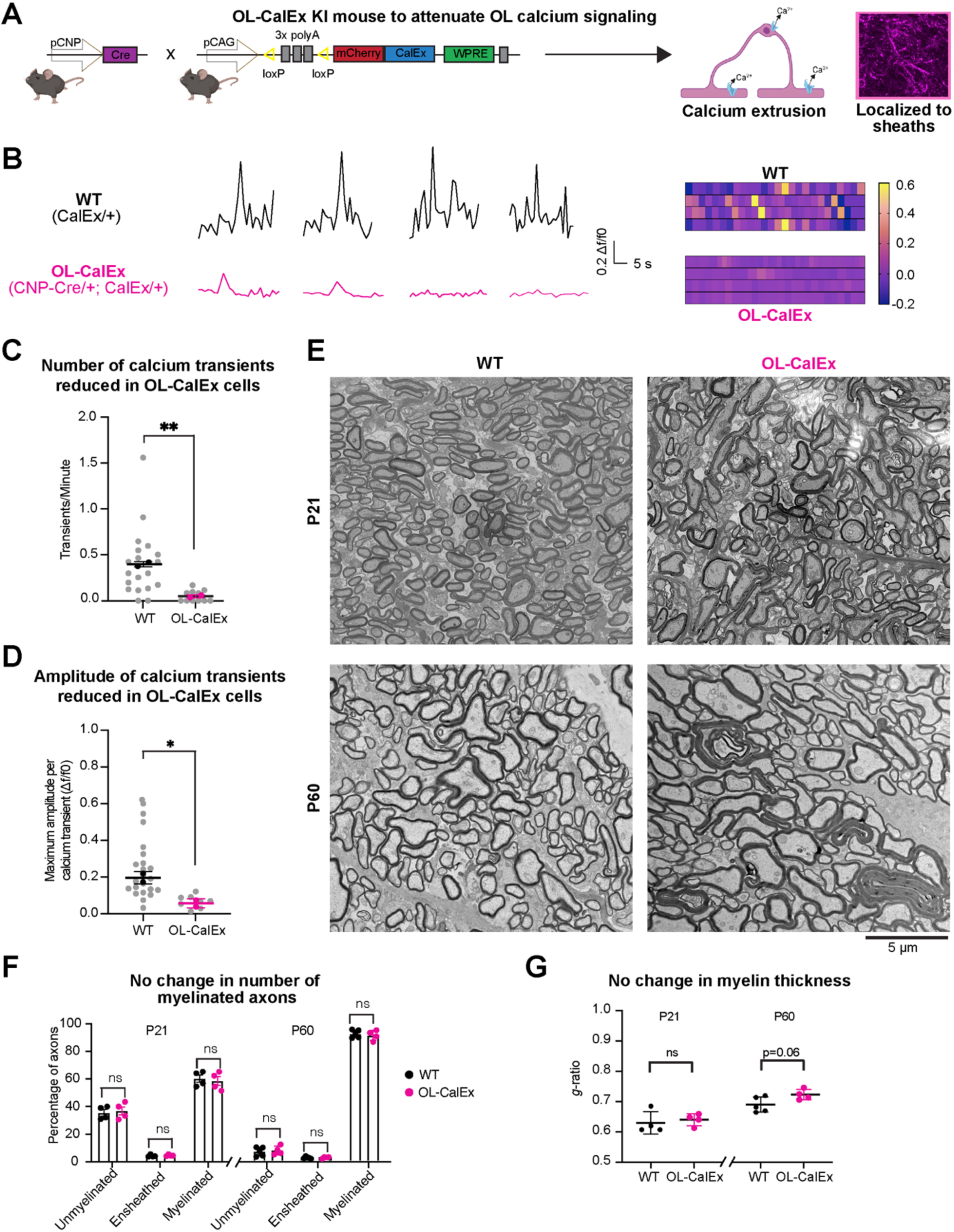
Attenuating oligodendrocyte calcium signaling does not grossly affect oligodendrocyte differentiation or myelination. (**A**) (left) Genetic strategy for attenuating calcium signaling in oligodendrocytes *in vivo*. CalEx^flox^ mice (floxed, transcriptional stop cassette in front of human plasma membrane calcium pump, hPMCA w/b) crossed to CNP-Cre. (right) Example of cortical oligodendrocyte expressing mCherry-CalEx. Created with Biorender.com. (**B**) Example calcium traces and corresponding heatmaps from (top) WT or (bottom) OL-CalEx primary oligodendrocytes. (**C**) Quantification of number of calcium transients per minute. Mean +/- SEM; N = 2 biological replicates (preps). Colored bolded dots represent average of individual biological replicates, grey dots represent values from individual cells. Statistical significance determined by unpaired, two-tailed Student’s t-test; **p < 0.01. (**D**) Quantification of calcium transient amplitude in WT and OL-CalEx oligodendrocytes. Average +/- SEM; N = 2 biological replicates (preps). Colored, bolded dots represent average of individual biological replicates, grey dots represent values from individual cells; *p < 0.05. (**E**) (top) Scanning Electron Microscopy of P21 mouse optic nerve cross sections (left: WT, right: OL-CalEx) (bottom) Transmission Electron Microscopy of P60 mouse optic nerve cross sections (left: WT, right: OL-CalEx), Scale bar, 5 μm. (**F**) Quantification of percentage of unmyelinated, ensheathed, and myelinated axons for P21 and P60 timepoints from electron microscopy in E. Average +/- SEM, N = 4 WT and N = 4 OL-CalEx for P21 and N = 5 WT and N = 4 OL-CalEx for P60. Statistical significance determined by multiple, unpaired, two-tailed Student’s t-test; n.s., not significant. (**G**) Quantification of myelin thickness via *g*-ratio for P21 and P60 timepoints from electron microscopy in E. Average +/- SEM, N = 4. Statistical significance determined by unpaired, two-tailed Student’s t-test; n.s., not significant.

We validated CalEx efficacy using calcium imaging of primary oligodendrocytes purified and cultured from OL-CalEx or WT littermates (Fig. 1, B-D, and Fig. S2G). In the absence of neurons, primary oligodendrocytes expand large, compact myelin membranes, making them a tractable model for studying oligodendrocyte cell biology^20, 35^. Primary cultured oligodendrocytes exhibit spontaneous calcium transients locally in their processes, even in the absence of neurons^36^. Compared to WT oligodendrocytes, CalEx-expressing oligodendrocytes had significantly fewer, lower-amplitude calcium transients (Fig. 1, B-D; Supplemental Video 1). CalEx expression did not affect oligodendrocyte survival, membrane expansion, or expression of MBP—a proxy for their differentiation and maturation into mature oligodendrocytes (Fig. S2A-F). Thus, CalEx is sufficient to attenuate calcium signaling in cultured oligodendrocytes without grossly perturbing their survival or differentiation.

We next quantified the specificity and penetrance of CalEx expression in oligodendrocytes in vivo in the developing mouse spinal cord during active myelination^15, 37^ (Fig. S1). As expected, CalEx expression was enriched in the white matter compared to neighboring grey matter (Figure S1C). On average, 93% of CalEx-expressing cells (mCherry+) were identifiably oligodendrocyte lineage cells (Olig2+; mCherry+). Of all CalEx-expressing OL-lineage cells, 75% were mature oligodendrocytes (CC1+; mCherry+), while the remaining 25% mCherry+ cells were premyelinating oligodendrocytes or oligodendrocyte precursor cells (OPCs) (CC1-/Olig2+; mCherry+) (Fig. S1E). 85% of all mature oligodendrocytes had detectable CalEx expression (Fig. S1F), indicating that CalEx expression is highly penetrant in these mice. Similar to our results showing that CalEx expression does not affect survival or differentiation and maturation in culture (Fig. S2A-F), the number of OPCs and differentiated oligodendrocytes was not significantly different between CalEx and WT littermates (Fig. S1G-I). Together, these results established CalEx as a precision tool for specifically attenuating oligodendrocyte calcium signaling during myelination.

### Oligodendrocyte calcium signaling regulates myelin membrane morphology during development

What is the consequence of calcium attenuation on myelination? Although calcium transients have been observed in nascent myelin sheaths in vivo^29–31^, it remains unknown whether they have any functional role in myelination. We analyzed myelin ultrastructure in CalEx and WT littermates using electron microscopy on optic nerves harvested during (P21) or at the end of (P60) developmental myelination (Fig. 1E-G). The optic nerve is ideal for quantifying myelination because its small size permits excellent preservation for ultrastructural studies, its axons are aligned, and the time course of myelination has been well-described^20, 38–40^. Surprisingly, CalEx expression had no effect on the number of myelinated axons at either timepoint (Fig. 1F). Myelin thickness, as quantified by *g*-ratio (the ratio between the inner and outer diameter of the myelin sheath), was also not significantly altered in OL-CalEx mice (Fig. 1G, Fig. S3C-D). Additionally, there were no differences in axonal caliber between the two groups at either timepoint (Fig. S3A-B).

Unexpectedly, OL-CalEx mice had a striking increase in the number of myelin outfoldings— improper outgrowths of myelin away from the axon (Fig. 2A-C). Outfoldings were frequently extremely long (length > twice the diameter of corresponding axon) and tortuous, extending into the neuropil where they often encircled neighboring myelinated axons or folded back onto themselves (Fig. 2A). We observed extremely-long outfoldings at both time points (P21 and P60), more commonly at P21 (Fig. 2, B-C).

**Figure 2.**
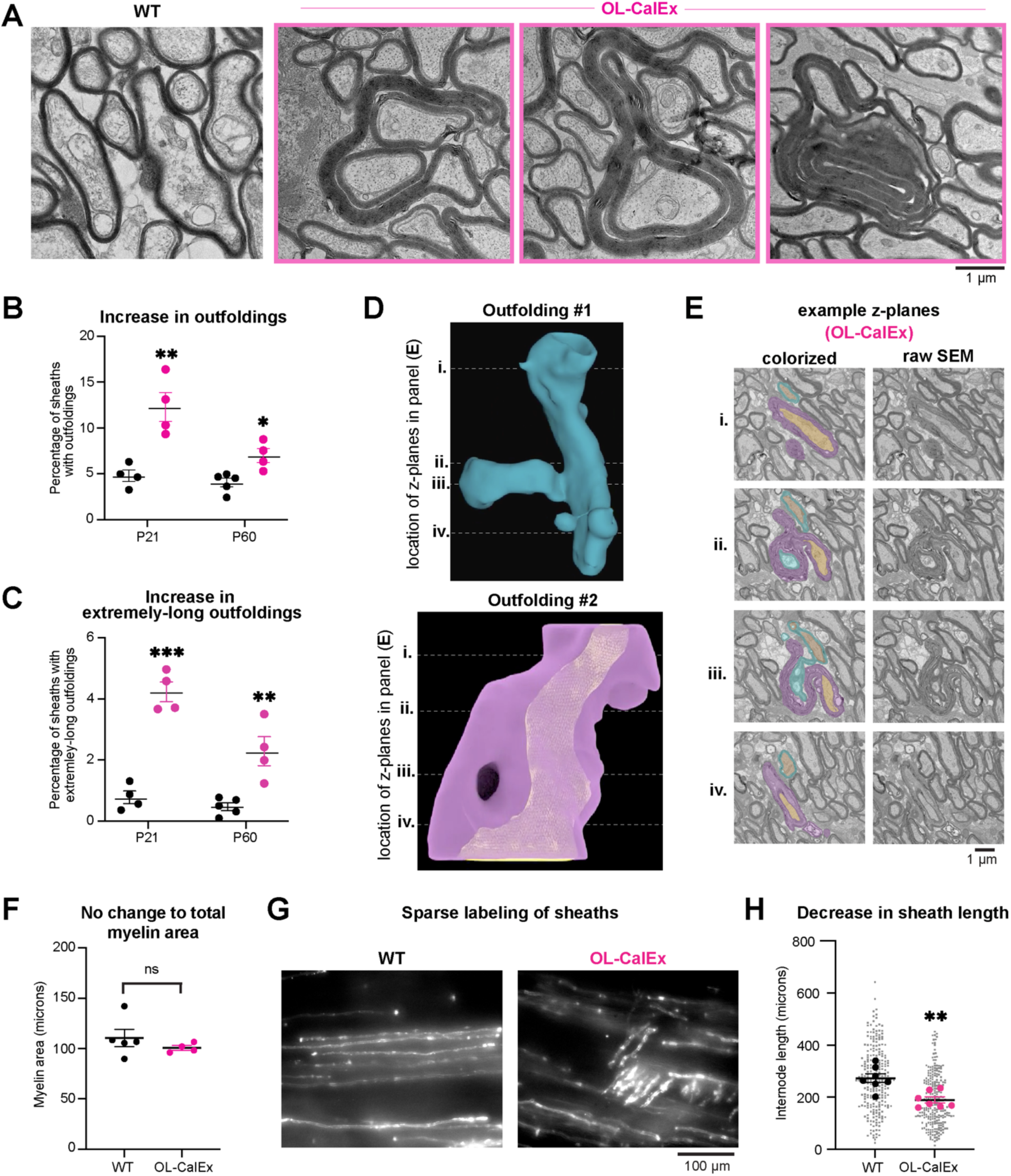
Oligodendrocyte calcium signaling is required for normal sheath length and myelin morphology. (**A**) Transmission electron microscopy (TEM) of WT (left micrograph) or OL-CalEx (right three micrographs) of P60 mouse optic nerve cross sections, highlighting examples of extremely-long outfoldings. Scale bar, 1 μm. (**B**) Quantification of percentages of myelin sheaths with outfoldings at P21 and P60 timepoints in (A). Average +/- SEM; P21, N = 4 WT and N = 4 OL-CalEx; P60, WT N = 5, OL-CalEx N = 4. Statistical significance determined by unpaired, two-tailed Student’s t-test; * p < 0.05, **p < 0.01. (**C**) Quantification of percentages of myelin sheaths with extremely-long outfoldings (defined as an outfolding greater than 2x the diameter of the axons) at P21 and P60 timepoints. Average +/- SEM; P21, N = 4; P60, WT N = 5, OL-CalEx N = 5. Statistical significance determined by unpaired, two-tailed Student’s t-test; **p < 0.01, ***p < 0.001. (**D**) 3-dimensional reconstruction of two proximal myelin outfoldings (top, blue; bottom, purple) and their respective axons (orange) from serial sectioning scanning electron microscopy (SEM). (**E**) Example (left) colorized and (right) raw SEM z planes through 3D reconstructions of both outfoldings. Dotted lines in (D) correspond to four numbered example z-planes through each outfolding. (**F**) Quantification of total myelin area from P60 TEM in Figure 1E. Average +/- SEM, WT N = 5, OL-CalEx N = 4; n.s., not significant. (**G**) Epifluorescence microscopy of P21 WT and OL-CalEx mouse whole-mount spinal cords with AAV-mediated sparse labeling of oligodendrocytes (myelin basic promoter driven EGFP). Scale bar, 100 μm. (**H**) Quantification of myelin sheath (internode) length from spinal cords in (G). Average +/- SEM, N = 7. Statistical significance determined by unpaired, two-tailed Student’s t-test; **p < 0.01

As an additional test of whether oligodendrocyte-autonomous calcium attenuation caused outfoldings, we employed an orthogonal genetic tool for blocking calcium signaling in cells—the genetic “calcium sponge” SpiCee^41^. SpiCee is a fusion protein containing four calcium-binding sites (two from calmodulin and two from a high-affinity parvalbumin variant) that attenuates cellular calcium signaling by binding and sequestering intracellular calcium, a mechanism distinct from CalEx (Fig. S4A)^41^. Compared to CalEx, the small size of SpiCee (∼615 bp) allows it to be packaged into adeno-associated virus (AAV) for in vivo delivery. We first validated that SpiCee-mRuby3 attenuated calcium signals in primary cultured oligodendrocytes, compared to control mRuby3 (Fig. S4, B and C). AAV-mediated expression of SpiCee in newly-formed oligodendrocytes in the developing spinal cord caused a ∼3-fold increase in the frequency of myelin outfoldings and a ∼20-fold increase in extremely-long outfoldings (Fig. S4, D-G). Combined with OL-CalEx experiments, these results suggested that outfoldings arise due to calcium attenuation specifically within oligodendrocytes.

To further examine the cellular basis of extremely-long outfoldings, we used serial electron microscopy to create 3D reconstructions of individual axon-myelin sheath pairs in OL-CalEx optic nerves (Fig. 2D-E; Supplemental Videos 2 and 3). 3D reconstructions revealed that individual myelin sheaths can have multiple outfoldings that originate from their exterior (abaxonal) sides, and outfoldings from neighboring sheaths can closely interact (Supplemental Video 2). We also observed outfoldings in OL-CalEx spinal cords at P8, indicating that outfoldings are not unique to the optic nerve and are formed by abnormal growth of myelin during the earliest stages of myelination (see Fig. 5, E-F). Myelin outfoldings were not accompanied by overall increase in myelin area (Fig. 2F), suggesting that myelin outfoldings represent inaccurate growth rather than overproduction of myelin.

Finally, given the lack of change in myelin area but the increase in outfoldings, we asked if the longitudinal extension of myelin sheaths along axons was affected in OL-CalEx. We used AAV-mediated sparse labeling of oligodendrocytes in the spinal cord to measure sheath length (internode length) at P21. Strikingly, OL-CalEx myelin sheaths were on average ∼30% shorter than wildtype, suggesting that calcium signaling in oligodendrocytes is required for the accurate longitudinal extension of myelin sheaths along axons (Fig. 2 G-H). Together, these data showed that oligodendrocyte calcium signaling is surprisingly dispensable for myelin initiation but is required instead to accurately sculpt the growth of myelin sheaths to achieve their normal length and morphology.

### Calcium signaling regulates actin filament levels in early-stage oligodendrocytes

How does attenuating calcium signaling in oligodendrocytes cause the myelination defects we observed? A major cellular target of calcium signaling is the actin cytoskeleton^42–46^, and actin dynamics are critical for myelination^19, 20^ Intriguingly, oligodendrocyte-specific deletion of proteins that promote actin filament assembly also cause abnormally long outfoldings, including the Arp2/3 complex that directly nucleates actin filaments necessary for early stages of myelination^20^ and N-Wasp^47^, a direct upstream activator of Arp2/3 (see Fig. 6). Thus, we speculated that outfoldings may arise in CalEx mice due to perturbed actin filament dynamics.

To test whether oligodendrocyte calcium signaling regulates actin, we first treated primary rat oligodendrocytes with cell-permeable calcium-chelating drugs (Fig. 3A) and measured their actin filament levels using phalloidin mean intensity in full-cell regions of interest (ROIs)^20^. Actin filament levels were significantly decreased by overnight treatment with either BAPTA (34% reduced) or dimethyl-BAPTA (DMB; 39% reduced) (Fig. 3B-C). Next, we purified primary mouse OPCs from OL-CalEx or WT littermates, induced their differentiation into oligodendrocytes, and visualized actin filaments (Fig. 3D). Similar to BAPTA/DMB treatment, actin filament levels were significantly decreased in mCherry-expressing OL-CalEx oligodendrocytes compared WT cells that do not express mCherry (24% reduced) (Fig. 3E-F). Oligodendrocytes in culture and in vivo have a “peak” of actin filament levels around the time when they are ensheathing axons^20^.

**Figure 3.**
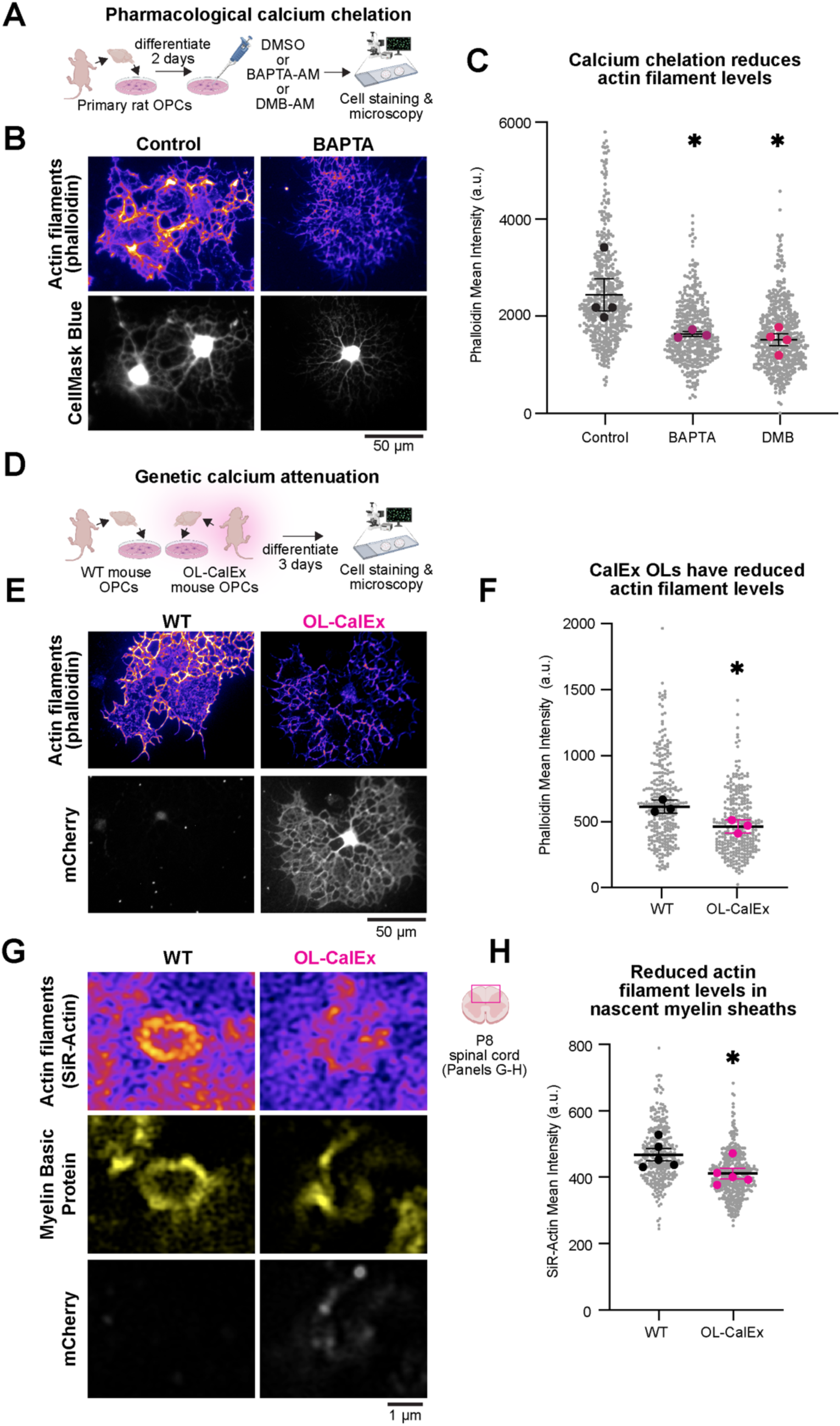
Calcium signaling in early stage oligodendrocytes regulates actin filament levels. (**A**) Primary rat oligodendrocyte precursor cells (OPCs) were differentiated for 2 days and then treated overnight with BAPTA-AM, dimethyl-BAPTA, or DMSO. Cells were then fixed, stained and imaged. Created with Biorender.com. (**B**) Epifluorescence representative micrograph of control (left) and BAPTA treated (right) rat oligodendrocytes stained with cell mask blue (white) to label cytoplasmic regions of the cells and phalloidin (purple) to label actin filaments. Scale bar, 50 μm. (**C**) Quantification of phalloidin mean intensity of rat oligodendrocytes in (B). Average +/- SEM, N = 4 biological replicates (preps/bolded dots) p-value was determined by a one-way ANOVA; *p < 0.05. (**D**) Oligodendrocyte precursor cells were isolated from WT or OL-CalEx mice and differentiated for 3 days before fixation and staining. Created with Biorender.com (**E**) Epifluorescence representative micrograph of WT (left) and OL-CalEx (right) mouse oligodendrocytes stained with cell mask blue (white) to label cytoplasmic regions of the cells and phalloidin (purple) to label actin filaments. Scale bar, 50 μm. (**F**) Quantification of phalloidin mean intensity of mouse oligodendrocytes in e. Average +/- SEM, N = 3 biological replicates (preps/bolded dots). Statistical significance determined by unpaired, two-tailed Student’s t-test; *p < 0.05. (**G**) Confocal micrograph of WT (left) and OL-CalEx (right) myelin rings in P8 dorsal spinal cord sections. (bottom row) mCherry staining (middle row) myelin basic protein to label nascent myelin sheaths (top row) SiR-Actin staining to label actin filaments. Scale bar, 1 μm. Created with Biorender.com. (**H**) Quantification of SiR-Actin mean intensity in myelin basic protein positive rings. Average +/- SEM, N = 5. Statistical significance determined by unpaired, two-tailed Student’s t-test; *p < 0.05

Intriguingly, oligodendrocyte actin was only sensitive to calcium attenuation at this exact stage of differentiation (Fig. S5G). As with CalEx expression (Fig. S2), treatment with BAPTA/DMB had no effect on oligodendrocyte differentiation, cell area, or cell survival for at least five days post-treatment (Fig. S5A-F). These results suggested that oligodendrocyte calcium signaling directly regulates actin, rather than indirectly by affecting differentiation.

Does calcium signaling also regulate the oligodendrocyte cytoskeleton *in vivo?* To look more closely at actin filaments in developing myelin sheaths, we co-stained spinal cord sections for myelin basic protein (MBP) and SiR-Actin, a cell-permeable, fluorogenic probe that specifically labels actin filaments^48^, and visualized individual sheaths using Airyscan (super-resolution) microscopy (Fig. 3G). Similar to our culture results, actin filament levels were decreased in (MBP+) nascent myelin sheaths (12% reduced; Fig. 3G-H). This figure may underestimate the actual reduction of actin filaments in nascent sheaths as the resolution limit of Airyscan microscopy (∼120 nm laterally) did not permit us to fully exclude SiR-Actin signal from axonal actin^49, 50^. Together, these data indicate that oligodendrocyte calcium signaling is required for normal actin filament levels in nascent myelin sheaths during development.

### Genetically inducing actin disassembly in oligodendrocytes phenocopies myelin morphology defects observed in OL-CalEx mice

Our results thus far predicted that myelin morphology defects may arise in OL-CalEx mice as a result of perturbed actin assembly in oligodendrocytes at the start of myelination. To test this idea, we used DeActs, genetically-encoded tools we developed to selectively induce actin disassembly in specific cell types in vivo^51^. We designed an oligodendrocyte-specific DeAct adeno-associated virus (AAV) construct using the myelin basic protein promoter (pMBP) to restrict DeAct expression to newly-formed and mature oligodendrocytes. To label DeAct-expressing myelin sheaths, this construct also expresses membrane-targeted EGFP (EGFP-caax) which is separated from DeAct-GS1 by a self-cleaving P2A sequence (Fig. 4A). We first confirmed that expression of pMBP-DeAct-GS1 induced actin disassembly in cultured OLs, while a control construct (pMBP-EGFP-caax alone) did not (Fig. S6A-B).

**Figure 4.**
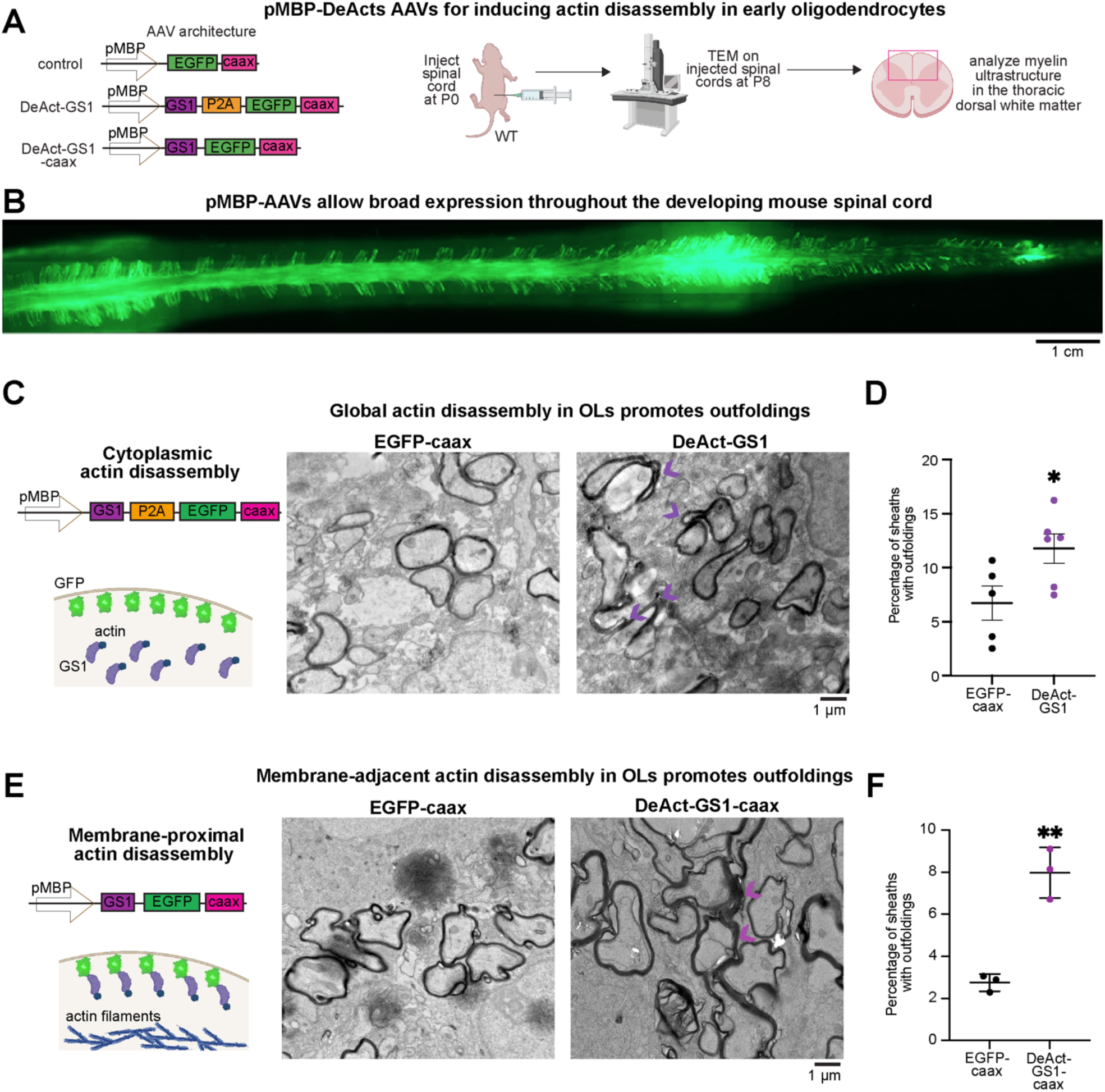
Genetically inducing actin disassembly in oligodendrocytes promotes myelin outfoldings. (**A**) (left) Control construct design: myelin basic protein promoter to drive expression in myelinating oligodendrocytes, followed by a farnesylated EGFP for visualization (EGFP-caax). Cytoplasmic DeAct (DeAct-GS1) construct design: myelin basic protein promoter, followed by Gs1 segment, followed by P2A self cleavable peptide, and a farnesylated EGFP for visualization. Membrane-targeted DeAct (DeAct-GS1-caax) construct design: myelin basic protein promoter, followed by a GS1 segment and farnesylated EGFP (one single fusion protein). (right) Virus encoding for DeAct-GS1 or DeAct-GS1-caax or control membrane targeted EGFP were injected into P0 mouse pups. Injected spinal cords were harvested at P8 and processed for transmission electron microscopy. Created with Biorender.com (**B**) Whole-mount spinal cord with EGFP expression after injection of AAV-pMBP-EGFP-caax. Scale bar, 1 cm. (**C**) Transmission Electron Microscopy of EGFP-caax injected (left) and DeAct-GS1 injected (right) spinal cord sections. Scale bar, 1 μm. (**D**) Quantification of percentage of myelin sheaths with outfoldings in (C). Average +/- SEM, N = 5 EGFP-caax, N = 6 DeAct-GS1. Statistical significance determined by unpaired, two-tailed Student’s t-test; *p < 0.05. (**E**) Transmission Electron Microscopy of membrane targeted EGFP injected (left) and DeAct-GS1-caax injected (right) spinal cord sections. Scale bar, 1 μm. (**F**) Quantification of percentage of myelin sheaths with outfoldings in d. Average +/- SEM, N = 3. Statistical significance determined by unpaired, two-tailed Student’s t-test; **p < 0.01.

To test whether inducing actin disassembly in early oligodendrocytes is sufficient to cause outfoldings, we targeted developing myelin in the mouse spinal cord. Compared to the inaccessible optic nerve, the dorsal spinal cord of neonatal mice is a tractable location for AAV injection and analysis of myelin ultrastructure^15, 20^. A single injection of pMBP-EGFP-caax AAV at P0 was sufficient to achieve bright, widespread EGFP expression in oligodendrocytes throughout the entire spinal cord by P8 and is specifically enriched in the white matter (Fig. 4B, S5C). Consistent with our prediction, inducing oligodendrocyte actin disassembly with DeAct-GS1 caused a 1.75-fold increase in myelin outfoldings (compared to EGFP-caax control), closely phenocopying OL-CalEx (Fig. 4C-D). Restricting DeAct-GS1 to the oligodendrocyte plasma membrane by direct fusion to a farnesyl-tag (DeAct-GS1-caax) also increased outfoldings (∼2.9 fold increase; Fig. 4E-F), suggesting that cortical actin—actin closely associated with the plasma membrane^52, 53^—may normally limit oligodendrocyte membrane expansion to prevent outfoldings (see Discussion).

### Genetically stabilizing actin in oligodendrocytes rescues myelin morphology defects in OL-CalEx mice

Would the stabilization of actin filaments in oligodendrocytes be sufficient to rescue the myelin morphology defects seen in the OL-CalEx mice? To test this idea, we developed a construct (Ezrin(abd*)-EGFP-caax) under the control of the MBP promoter that causes an oligodendrocyte specific overexpression of the constitutively-active actin binding domain of the cortical actin-binding protein Ezrin, which has been previously used in other cell types to induce stabilization of cortical actin (Fig. 5A)^52^. We first confirmed that the expression of the Ezrin(abd*)-EGFP-caax increased actin filament levels in cultured oligodendrocytes compared to an EGFP-caax control (Fig. 5B-C).

**Figure 5.**
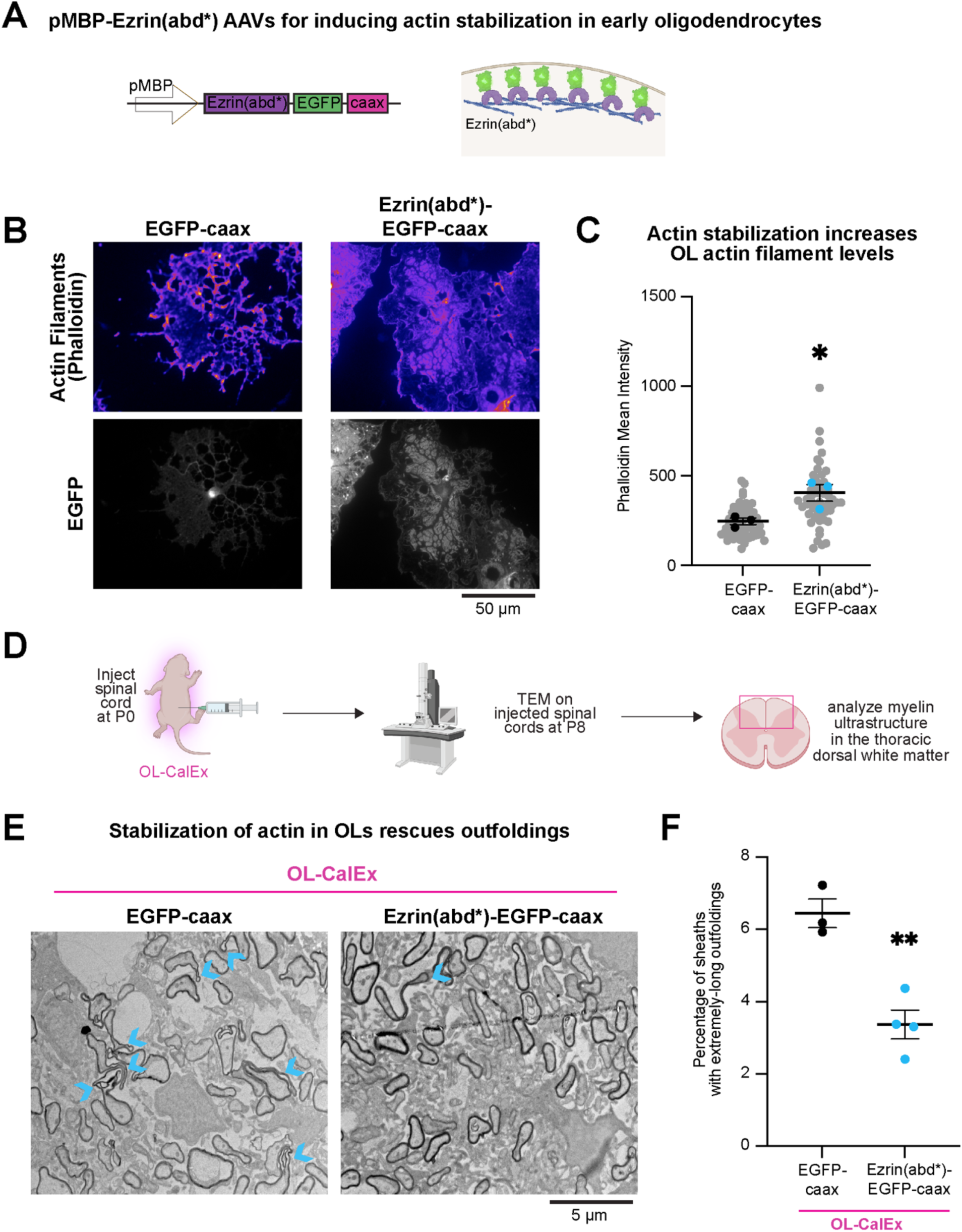
Genetically inducing actin stabilization in CalEx oligodendrocytes rescues myelin outfoldings. (**A**) Construct design to stabilize membrane-proximal actin in developing myelin sheaths: myelin basic promoter (pMBP), followed by constitutively-active Ezrin actin binding domain (Ezrin(abd*)) fused to EGFP-caax. Created with Biorender.com. (**B**) Epifluorescence representative micrograph of EGFP-caax (left) and Ezrin(abd*)-EGFP-caax (right) transfected oligodendrocytes stained with phalloidin (purple) to label actin filaments (top row) and endogenous EGFP expression (grey, bottom row). Scale bar, 50 μm. (**C**) Quantification of phalloidin mean intensity rat oligodendrocytes in b. Average +/- SEM, N = 3 biological replicates (preps/bolded dots). Statistical significance determined by unpaired, two-tailed Student’s t-test; *p < 0.05 (**D**) Virus encoding for Ezrin(abd*)-EGFP-caax or control EGFP-caax were injected into P0 mouse pups. Injected spinal cords were harvested at P8 and processed for transmission electron microscopy. Created with Biorender.com (**E**) Transmission Electron Microscopy of EGFP-caax injected (left) and Ezrin(abd*)-EGFP-caax (right) spinal cord sections. Scale bar, 5 μm. (**F**) Quantification of percentage of myelin sheaths with extremely-long outfoldings in (E). Average +/- SEM, N = 3. Statistical significance determined by unpaired, two-tailed Student’s t-test; *p < 0.05, **p < 0.01.

To test whether stabilization of actin filaments in oligodendrocytes is sufficient to rescue the outfolding defect in OL-CalEx mice, we injected AAVs encoding for either MBP promoter driven EGFP-caax or Ezrn(abd*)-EGFP-caax into OL-CalEx pups (Fig. 5D). Consistent with the prediction that stabilizing actin filaments in developing myelin sheaths would rescue the outfolding defect, inducing actin stabilization in oligodendrocytes caused a 1.9-fold reduction in outfoldings, indicating that actin filament restoration is partially capable of restoring calcium-attenuation defects (Fig. 5E-F). Together, these results indicate that oligodendrocyte calcium signaling is required for actin-dependent regulation of myelin membrane morphology—and raise the possibility that this mechanism may allow myelin sheaths to respond to neuronal properties including activity to fine-tune conduction velocity (Fig. 6).

**Figure 6.**
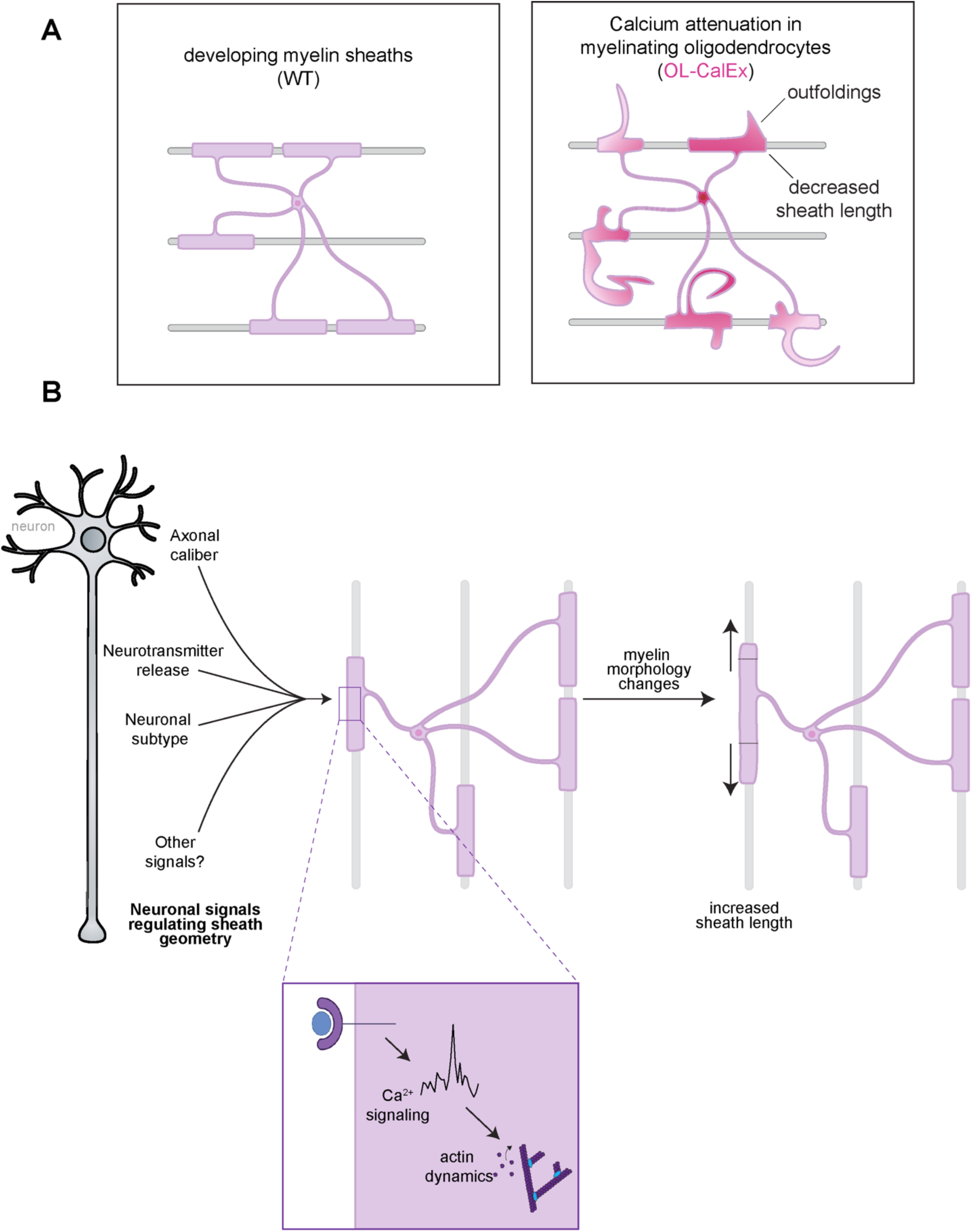
Model figure for oligodendrocyte calcium regulation of myelin sheath morphology. (**A**) (left) In developing myelin sheaths, calcium regulates actin filaments, which are required to guide myelin membrane accurately around neuronal axons. (right) Loss of calcium signaling leads to extremely-long myelin outfoldings, which are caused by mistargeted myelin. (**B**) Model for how neuronal activity may regulate activity dependent myelination through calcium signaling in oligodendrocytes. Active neurons release neurotransmitters and other to-be-identified factors that can bind to receptors on oligodendrocytes. This receptor-ligand binding could modulate oligodendrocyte calcium signaling, which in turn could regulate actin dynamics. These cell-biological changes in oligodendrocytes could therefore regulate changes to myelin morphology.

## DISCUSSION

How myelin sheaths are precisely sculpted during development and remodeled in adults to fine-tune nerve conduction velocity are fundamental questions in neurobiology. Prior studies suggest that calcium signaling in myelin sheaths is a potential mechanism that could allow oligodendrocytes to shape their myelin sheaths in response to the demands of the neurons they myelinate^29–31^. Here, we tested this idea by genetically attenuating calcium signaling in newly-formed oligodendrocytes in the developing mouse CNS using the genetic calcium pump CalEx. We found that calcium signaling is surprisingly dispensable for the differentiation and maturation of oligodendrocytes or for the initiation of myelination. However, attenuation of oligodendrocyte calcium signaling leads to shorter myelin sheaths and long outfoldings, suggesting that the major role of local calcium signaling in myelin sheaths is to properly sculpt sheath morphology. Mechanistically, we found that calcium signaling promotes actin cytoskeletal assembly in nascent sheaths. Accordingly, genetically inducing actin disassembly phenocopied the loss of calcium signaling in oligodendrocytes, while stabilization of actin filaments partially rescued the myelin outfolding defect in CalEx mice. Thus, a major role of calcium signaling in oligodendrocytes during developmental myelination is actin-dependent regulation of myelin morphology.

Oligodendrocyte calcium signaling may be a mechanism by which an oligodendrocyte can adjust its myelination patterns based on properties of the neuron that it is myelinating. Previous work has shown that myelin sheath length and thickness are dictated by neuronal properties including axonal caliber, neuronal type, and neuronal activity^10^. However, how myelin senses and responds to neuronal properties to build sheaths of the proper length and thickness remain unknown. Calcium influx into the cytosol can be regulated by a diverse set of mechanisms including voltage gated and mechanosensitive channels, store operated channels and release from the endoplasmic reticulum, mitochondrial permeability pore opening, sodium/calcium exchangers, gap junctions, and neurotransmitter receptors—all of which may regulate calcium signaling in oligodendrocytes in response to the properties of the neurons being myelinated. For example, calcium signaling in oligodendrocytes can be modulated by neuronal activity in zebrafish^30^, and can be triggered by neurotransmitters including ATP and glutamate—at least in cultured oligodendrocytes^27, 28^. Oligodendrocyte calcium signaling is thus poised to act as an essential cell-signaling bridge between these various neuronal properties and myelin.

How does loss of calcium signaling cause myelin outfoldings? One possibility is that the outfoldings seen in OL-CalEx mice are a direct consequence of having shorter sheaths—the same amount of myelin membrane extending out into the neuropil rather than stretching along the axon (by analogy, outfoldings similar to the folds of a scrunched-down sock). If true, the pathological presence of outfoldings observed in other myelin mutants and neuropathies^11–14, 17, 20, 54, 55^ may also be a consequence of improper sheath extension. However, it remains possible that outfolding formation occurs independent of sheath length. In other cell types, cortical actin—actin closely associated with the plasma membrane—limits membrane expansion^52, 53^. Our experiments using membrane-tethered tools to induce actin filament disassembly (Fig. 4) or stabilize actin filaments (Fig. 5) at the plasma membrane suggest that a membrane-proximal actin cytoskeleton could locally restrict aberrant myelin membrane extension away from sheaths. This model predicts that regulated, local loss of the actin cytoskeleton—e.g. in the oligodendrocyte “inner tongue” (the leading edge that grows around the axon during wrapping)—could allow for myelin growth to be specifically confined to spirally wrapping around the axon, rather than away from the axon as in an outfolding. Whether newly differentiated oligodendrocytes have membrane-adjacent actin subpopulations and the role that these actin populations play in regulating myelin membrane growth remain open questions. As a final potential explanation for the cause of outfoldings, it was recently found that outfoldings are present during early development and then resolve over time, and may require “pruning” by microglia to fully resolve^10, 17, 56, 57^. It is possible that attenuating oligodendrocyte calcium signaling perturbs this normal process of outfolding resolution, perhaps by inhibiting microglial pruning.

Actin filament dynamics are critical for normal myelin formation in the CNS^19, 20^. We previously found that distinct stages of myelination have opposite requirements for actin filaments: actin filament assembly is required for oligodendrocytes to extend their cellular processes to ensheath axons, but the subsequent stage of myelin wrapping requires actin filaments to disassemble^20^. Our findings using CalEx mice agree with and build from this earlier work. Calcium signaling promotes actin filament assembly in nascent sheaths/early-stage oligodendrocytes, but is not necessary for actin disassembly at later stages (Fig. S5). Accordingly, myelin wrapping—which requires actin disassembly—is unaffected by calcium attenuation (Fig. 1).

What are the molecular mechanisms that control calcium-mediated actin assembly in oligodendrocytes? One possible mechanism is via gelsolin, an actin disassembly factor that is highly expressed by oligodendrocytes^20, 58^ and directly activated by calcium^59^. Inconsistent with gelsolin mediating the effect of calcium on actin regulation in oligodendrocytes, gelsolin knockout mice show no evidence of outfoldings (Fig. S7A-C)^20^ and the major role of gelsolin during myelination is to induce actin disassembly to drive wrapping^20^. Therefore, it is unlikely that gelsolin directly mediates calcium’s control over actin assembly in early stages of myelination. A second model for how calcium may regulate actin in oligodendrocytes is via its established role in promoting actin assembly through phosphoinositide (PI_4,5_P_2_) signaling and the N-Wasp-Arp2/3 axis^17^. Intriguingly, prior work^20, 47, 54, 55^ and our findings here show that interfering with any step in this signaling pathway causes myelin outfoldings (Fig. S7D). Together, these results converge on a model for myelin outfolding formation that may be shared by many disease states and myelin mutants—dysregulated actin assembly. It will be interesting to test this model in future studies.

What other cell biological processes does calcium regulate in oligodendrocytes? Recently, we discovered that myelin membrane is added via SNARE-mediated exocytosis^15^, a process that is well-known to be regulated by calcium in many other cell types^25, 60, 61^. Future studies will focus on whether oligodendrocyte exocytosis is regulated by calcium, and how actin and exocytosis may collaborate^62^ to regulate the morphology of myelin sheaths. In contrast to the importance of calcium signaling during myelin formation, dysregulation of oligodendrocyte calcium signaling may be pathogenic in disease states^28^. Forced calcium influx into mature sheaths is sufficient to cause myelin decompaction and breakdown, a form of myelin pathology seen in response to numerous demyelinating insults^63, 64^. Thus, it would be interesting to test whether CalEx-mediated calcium attenuation protects against demyelination in these models.

Does oligodendrocyte calcium signaling also drive myelin remodeling to support learning? Experience and neuronal activity cause dramatic changes to the pattern of myelin in the adult brain, including addition of new myelin sheaths and morphological changes to existing sheaths^3–9^. Although such changes are predicted to affect the function of neuronal circuits, the overarching role of myelin dynamics in learning is still mostly hypothetical. An essential next step for the field is to determine the cellular mechanisms that drive activity-induced myelin remodeling, and then leverage that knowledge to be able to test the importance of myelin remodeling across learning modalities. Could oligodendrocyte calcium signaling be the missing mechanistic link between neuronal activity and plasticity-induced changes to myelin? While the precise role of neuronal activity in driving oligodendrocyte calcium signaling remains an open question^31^, several lines of evidence converge to support the idea that neuronal activity could regulate oligodendrocyte sheath morphology via calcium signaling. First, blocking neuronal activity or neurotransmitter release perturbs myelin sheath lengths in zebrafish^65–68^. Second, several neurotransmitters including glutamate and ATP are sufficient to elicit calcium transients in oligodendrocytes, although the extent to which this effect persists as oligodendrocytes mature remains to be elucidated^27, 28^. Third, neuronal activity promotes sheath elongation^65^ and calcium transients in sheaths^30^. Together, these results suggest that local calcium transients in sheaths could respond to neuronal activity to control sheath remodeling. Although these findings were specific to development, similar mechanisms could potentially occur during learning-induced oligodendrogenesis or sheath remodeling in the adult. For example, it is possible that neuronal activity-induced calcium signaling in oligodendrocytes directly regulates actin dynamics to power plasticity-induced myelin remodeling, similar to calcium’s role in nascent sheaths during development that we uncovered here.

In summary, we discovered that calcium signaling is required for the actin-dependent sculpting of myelin sheath morphology during development. We propose that this mechanism allows myelin sheaths to tune conduction velocity in response to diverse neuronal signals, including activity. Our work provides a conceptual framework for future studies to understand how oligodendrocyte calcium signaling contributes to myelin remodeling during learning and myelin loss/regeneration in the context of disease.

## ACKNOWLEDGMENTS

We thank current and past members of the Zuchero lab (especially the OPC Crew: Madeline Cooper, Maya Weigel, Graham Jones, and Miguel Garcia), Michael Burks, and Ganesh and Neeraja Iyer for their helpful discussions and support. We also thank Ethan Hughes for providing insightful feedback on the manuscript. We thank Klaus Nave for kindly sharing the *Cnp-Cre* mouse line. We also thank the Stanford University Cell Sciences Imaging Core Facility for Transmission Electron Microscopy data collection, especially John Perrino and Ibanri Phanwar-Wood for their expertise in processing and staining EM samples (RRID:SCR_017787: supported by an ARRA Award Number 1S10RR026780-01 from the National Center for Research Resources. Its contents are solely the responsibility of the authors and do not necessarily represent the official views of the NCRR or the National Institutes of Health.). Electron microscopy image processing was supported in part by the grants: NN1 NSF 1707356 and NN2 NSF 2014862. The authors acknowledge the Texas Advanced Computing Center (TACC) at The University of Texas at Austin for providing HPC and visualization resources that have contributed to the research results reported within this paper. We also thank the Stanford Neuroscience Gene Vector and Virus Core for producing the adeno-associated viruses used in this study. Images and diagrams were created using BioRender.com.

This project was supported by the Regina Casper Stanford Graduate Fellowship (M.I.), the Stanford Berry Postdoctoral Fellowship (H.K.), Waitt Foundation and Core Grant application NCI CCSG (CA014195) (S.W.N, L.R.A, U.M.), Helen Hay Whitney Foundation (M.L.), Stanford Wu Tsai Neurosciences Interdisciplinary Scholar Award (M.L.), NINDS R35NS111583 (X.Y.,B.S.K.), Chan-Zuckerberg Imaging Scientist Grant (U.M), NSF Neuronex Grant 2014862 (U.M.), the McKnight Endowment Fund for Neuroscience (J.B.Z.), the Stanford Bio-X Interdisciplinary Initiatives Seed Grants Program (IIP) [R9-24] (J.B.Z.), the National Multiple Sclerosis Society Harry Weaver Neuroscience Scholar Award (J.B.Z.), the Beckman Young Investigator Award (J.B.Z.), the Myra Reinhard Family Foundation (J.B.Z.), the National Institutes of Health R01NS119823 (J.B.Z.), and the Koret Family Foundation (J.B.Z.).

## AUTHOR CONTRIBUTIONS

MI and JBZ conceived of the project. MI designed, performed, and analyzed all experiments with the following exceptions. HK and NA performed DeAct-Gs1 spinal cord injections and electron microscopy in Figure 4. SWN and LA performed SEM imaging and analysis on P21 samples in Figure 1 and 3DSEM reconstructions in Figure 2 with guidance from UM. ML optimized characterization pipeline used in used in Figure S1. XY and BSK developed and validated CalEx^flox^ line used in the present study. MI and JBZ wrote the manuscript. All authors provided feedback on the manuscript.

## STAR*METHODS

### Animals

All procedures involving animals were approved by the Institutional Administrative Panel on Laboratory Animal Care (APLAC) of Stanford University and followed the National Institutes of Health guidelines. Animals were group-housed in a Stanford University animal facility with a 12:12 hour light/dark cycle and ad libitum access to water and food. All mice were monitored by veterinary and animal care staff. Animals used in the study had not undergone prior procedures. CalEx fl/fl (which contain a floxed STOP cassette in front of CalEx-mCherry) mice were gifted by Drs. Xinzhu Yu and Baljit Khakh. The CNP-Cre mice were gifted by Dr. Klaus Armin-Nave. The OL-CalEx line was generated by crossing homozygote CalEx fl/fl with heterozygote CNP-Cre/+ mice. The following primers were used for genotyping: Common CalEx forward primer 5’– CCTTTCTGGGAGTTCTCTGCTGC–3’; WT Reverse CalEx 5’– GCGGATCACAAGCAATAATAACCTG–3’ Mutant Reverse CalEx primer 5’– CGTAAGTTATGTAACGCGGAACTCC–3’. CNP forward primer 5’—GCCTTCAAACTGTCCATCTC—3’; CNP reverse 5’—CACCATTATTTTCCCGACCC—3’. All resulting progeny from these crosses were used; CalEx/+; CNP-Cre/+ mice are referred to as “OL-CalEx.” Sprague-Dawley rats and C57BL/6 mice were ordered from Charles River Laboratories. Male and female mice were used for all *in vivo* experiments. For cell culture studies, brains of both sexes were pooled to obtain sufficient cell numbers.

For all histology, animals were anesthetized with injecting a cocktail of ketamine (100 mg/kg) and xylazine (20 mg/kg). Animals were then transcardially perfused with PBS and tissues were dissected and drop fixed with 4% paraformaldehyde (PFA) (made from 16% PFA, Electron Microscopy Sciences) for immunohistochemistry or Karlsson and Schultz (KS) fix for transmission electron microscopy.

### Isolation of Oligodendrocyte Precursor Cells

Primary oligodendrocyte precursors were purified by immunopanning from P5-P7 Sprague Dawley rat or P6-P8 transgenic mouse brains as previously described. Oligodendrocyte precursors were typically seeded at a density of 50,000-250,000 cells/10-cm dish and recovered for 4 days in culture before lifting cells via trypsinization and distributing for transfection, proliferation, or differentiation assays. All plasticware for culturing oligodendrocyte precursors were coated with 0.01 mg/ml poly-D-lysine hydrobromide (PDL, Sigma P6407) resuspended in water. All glass coverslips for culturing oligodendrocyte precursors were coated with 0.01 mg/ml PDL, which was first resuspended at 100x in 150 mM boric acid pH 8.4 (PDL borate).

To proliferate primary oligodendrocyte precursors, cells were cultured in a serum-free defined media (or “DMEM-SATO” base media) supplemented with 4.2 μg/ml forskolin (Sigma-Aldrich,Cat#F6886), 10 ng/ml PDGF (Peprotech, Cat#100-13A), 10 ng/ml CNTF (Peprotech, Cat#450-02), and 1 ng/ml neurotrophin-3 (NT-3; Peprotech, Cat#450-03) and house in an incubator set to 37C with 10% CO2. To induce differentiation, cells were switched to DMEM-SATO base media containing 4.2 μg/ml forskolin (Sigma-Aldrich, Cat#F6886), 10 ng/ml CNTF (Peprotech, Cat#450-02), 40 ng/ml thyroid hormone (T3; Sigma-Aldrich, Cat#T6397), and 1x NS21-MAX (R&D Systems AR008).

### Plating Schemes and Pharmacological Cell Treatments

Primary oligodendrocyte precursors from CalEx;Cnp-Cre and control littermate mice were harvested and cultured as described above in “Purification and Culturing of Cells”. Cells were seeded onto 12-mm glass coverslips (Carolina Biological Supply No. 63-3029) at a density of 5,000 cells/coverslip in differentiation media. Cells were cultured until day three in differentiation and then fixed for subsequent staining analysis.

For all BAPTA-AM (ThermoFisher B6769) or dimethyl BAPTA-AM (Sigma-Aldrich 16609-10MG-F) experiments, wildtype rat oligodendrocyte precursors were plated in differentiation media at a density of 10,000 cells/coverslip. Briefly, BAPTA-AM or dimethyl-BAPTA-AM stocks were resuspended in tissue culture grade DMSO at a concentration of 1uM. Drugs were then diluted to 2x final concentration in completed DMEM-SATO with differentiation factors. 250 uL of media was removed from each well and replaced with 250 uL of drug or DMSO containing media. Final drug concentrations were 1uM or 500 nM. Previous reports have found that at high micromolar concentrations, BAPTA can cause calcium-independent disassembly of the cytoskeleton in cultured cells, but that variants of BAPTA such as di-methyl BAPTA do not have these off-target effects on the cytoskeleton ^69^.

### Design of DeAct-GS1-Caax and Ezrin(ABD*) constructs

DeAct-GS1-Caax and Ezrin(ABD*) constructs were created using InFusion cloning (Takara Bio) by annealing one DNA fragment into the parent plasmid. The parent plasmid contains a 1.3kb MBP promoter driving expression of (farnesylated, membrane-bound) EGFP-caax in a pAAV vector backbone and was linearized using NcoI and BglI. The fragment encoding for either GS1 (aa 53-176, Addgene # 89445; Reference 56) or Ezrin(ABD*) (Ezrin aa 552-586 with T567D constitutively-active mutation. Addgene #155227) was cloned with 15 base pair overhangs required for InFusion reactions. This created constructs with the following configuration: pMBP-GS1-EGFP-caax or Ezrin(ABD*)-EGFP-caax.

### Transfection of oligodendrocyte precursors

Rat oligodendrocyte precursor cells were trypsinized and dissociated from tissue culture dishes and centrifuged at 700xrpm for 10 min. 250,000 oligodendrocyte precursors were gently resuspended in 20 μl of nucleofector solution (Lonza P3 Primary Cell 4D-Nucleofector V4XP-3032) with 400 ng of GFP-Caax, Gs1-P2a-GFP Caax, or Gs1-Caax plasmids. Cells were loaded into a 16-well cuvette and electroporated in a Lonza 4D-Nucleofector X Unit (AAF-1003X) assembled with a 4D-Nucleofector Core Unit (AAF-1002B) using pulse code DC-218.

Electroporated cells rested for 10 min at room temperature. 80 μl of antibiotic free DMEM-SATO media was added to each cuvette and cells were gently triturated. Each well of 250k cells was distributed onto No. 1 glass coverslips coated with PDL-borate for differentiation timepoints and technical replicates (up to six coverslips from a single transfection). Each coverslip was half-fed with freshly supplemented DMEM-SATO media every two to three days.

### Calcium Imaging

Immunopanned CalEx; Cnp-Cre oligodendrocyte precursors were differentiated for two-three days for all calcium imaging experiments. To load 1 uM cell permeable calcium indicator, Fluo-4-AM (Invitrogen, F14201) 50 µg tube of lyophilized Fluo-4-AM stock was resuspended in 46 uL of sterile DMSO (1mM concentration). 1.5 uL of Fluo-4-AM stock was added directly to cell media to prevent shearing of cells. Cells were incubated with Fluo for 20 minutes and then media was completely changed to 1 mL DMEM-SATO Media (made with Fisher Scientific A1896701) for imaging.

Cells were imaged on an Opterra II Multipoint Swept Field Confocal outfitted with a humidified, temperature-controlled microscope enclosure (Okolab microscope enclosure, H201-Temperature Unit, CO2 controller, HM-Active Vibration Free Humidity Controller with humidity sensor and temperature-controlled tube). Imaging was performed using the 60x/1.2 NA water objective and Perfect Focus to prevent z-plane drift during imaging. Images were acquired every two seconds in 488 channel with 70 uM slit and 100 ms exposure time with 15% laser power. Calcium imaging data was analyzed using Fiji to select regions of interest and was analyzed to extract rate and amplitude measurements. Data was analyzed blind to condition.

### Primary Oligodendrocyte Antibody Staining

At the specified day of differentiation, cell media was removed, and coverslips were treated with 4% PFA for 15 min at RT, followed by three washes with 1xPBS and permeabilization in 0.1% Triton X-100 in PBS for 3 min at RT. Prior to staining, cells were incubated in a blocking solution of 3% BSA in PBS for 20 min at RT. Then, primary antibodies (rat anti-MBP and/or rabbit anti-RFP) were added in a 3% BSA solution for overnight incubation at 4C. On the following day, the primary antibody solution was rinsed off with three washes of PBS, and then incubated with secondary antibodies (anti-rat AlexaFluor 594 or 647) in 3% BSA for 1 hr at RT. After three washes with PBS, CellMask Blue stain (1:1000) and phalloidin-488 (7 uL phalloidin per 1 mL PBS) was incubated to stain all cells for 15 min at RT, followed by three additional rounds of washing with PBS. Stained cells were mounted onto microscope slides (Fisher Scientific 12-550-143) in Fluoromount G (SouthernBioTech, 0100-20).

Cells were imaged by widefield epifluorescence with a Zeiss Axio Observer Z1 using the Plan-Apo 20x/0.8 NA objective for actin, MBP and cell area quantifications. Images were acquired blinded to the genotype or condition with identical illumination and acquisition conditions per biological replicate.

### Spinal Cord Injections

AAV-DJ serotype virus was produced by the Gene Vector Virus Core at Stanford University and stored at −80°C until use in experiments. Virus was thawed and diluted 1:1 with Trypan Blue for high-titer injections or 1:8 (for 1 uL of virus, 1 uL of Trypan Blue and 6 uL of sterile D-PBS) for sparse labeling experiments. P0-P1 C57/Bl6 mouse pups were placed in a KimWipe and deeply anesthetized on ice until unresponsive to a toepinch. Using a Hamilton syringe (Model 80308 701SN, Point Style 4, 32 gauge, 20 mm length, and 12°), AAVs were injected into the lumbar spinal cord of mouse pups; successful injections lead to a blue “stripe,” indicating that virus had traveled throughout the spinal cord. Pups recovered at 34°C on a heating pad until pink and moving freely and then were returned to the home cage. Spinal cords were then extracted as described (see “Animals”) and processed either for IHC or TEM.

### Spinal Cord Immunofluorescent Staining

Freshly dissected spinal cords were drop fixed in 4% paraformaldehyde for 5 hours, washed with PBS, and then dropped into 30% sucrose overnight. Fixed spinal cords were sectioned on a cryostat (CM3050S, Leica Microsystems) into 30 μm sections (Olig2/CC1 staining) or 10 μm sections (SiRActin/MBP staining), and immediately mounted onto Superfrost Plus (VWR) microscope slides. Slides were dried at 34°C for 10 minutes to ensure section adherence, and then stored at −80°C until subsequent use.

Prior to staining, slides were warmed at 34°C for 30 minutes, sections were circled with a hydrophobic pen (ThermoScientific 008899) and dried for another 20 minutes. Sections were permeabilized with PBST (0.1% Triton X-100 in PBS) for 5 minutes at RT, blocked with 10% donkey serum in PBST for 1 hour and then primary antibodies diluted in 1% donkey serum in PBST were applied (dilutions specified in “Antibodies” section) and incubated at 4°C overnight. The next day, primary antibody was rinsed with 3 x 20 minute washes in PBS. Secondary antibodies and SiRActin diluted in 1% donkey serum in PBST were incubated for 2 hours at RT. Secondary antibody was washed 3 x 20 minutes in PBS. Samples were mounted in Vectashield with DAPI (Vector, H-1200) using a coverslip.

### Antibodies and Staining Reagents

Primary antibodies used in this study were as follows: Rt-anti-MBP (Abcam ab7349; 1:100), Ms-anti-CC1 (1:500), Goat-anti-Olig2 (Millipore AB9610; 1:500), Rb-anti-RFP (Rockland #600-401-379; 1:1000).

Secondary antibodies and reagents conjugated to fluorescent dyes used in this study were as follows at a 1:1000 dilution for both tissue and primary cell immunofluorescence: donkey anti-rat Alexa Fluor 594 (Thermo Scientific A-21209), goat anti-rat Alexa Fluor 647 (Thermo Scientific A-21247), donkey anti-mouse Alexa Fluor 488 (Thermo Scientific A-21202), donkey anti-mouse Alexa Fluor 647 (Thermo Scientific A-31571), donkey anti-rabbit Alexa Fluor 594 (Thermo Scientific, A-21207) Alexa Fluor 488 conjugated Phalloidin (Thermo Scientfic, A12379), HCS CellMask Blue (Thermo Scientific H32720).

### SEM sample preparation

Samples were processed and imaged as previously described^70^. Materials were sourced from Electron Microscopy Sciences (Hatfield, PA) unless otherwise stated. Samples were processed and imaged as described elsewhere. Fixed samples of optic nerve were carefully dissected and immersed in a modified Karnovsky’s fixative (4% paraformaldehyde, 2.5% glutaraldehyde, 0.1M sodium cacodylate, 3mM calcium chloride) for at least 24 hours. Samples were rinsed repeatedly with ice-cold buffer (0.1M sodium cacodylate, 3mM calcium chloride) before further fixation with reduced osmium (1% osmium tetroxide, 1.5% potassium ferrocyanide, 0.1M sodium cacodylate, 3mM calcium chloride). Samples were rinsed repeatedly with ice cold water and stained with 1% aqueous uranyl acetate overnight at 4°C. The following day, samples were rinsed thoroughly with ice cold water and serially dehydrated in ascending concentrations of ice cold ethanol. Samples were finally rinsed in three changes of anhydrous ethanol at room temperature before infiltration with a 1:1 mixture of anhydrous ethanol and epoxy resin (Epon 812, hard formulation) for 4 hours on a rotating mixer, followed by infiltration in pure resin overnight on a rotating mixer.

The following morning, samples were transferred into polypropylene bottlecaps with fresh resin and allowed to infiltrate for two hours before flat embedding in silicone molds, with nerves oriented to facilitate cross-sectioning. Blocks were polymerized at 70°C for 48 hours.

Ultra thin sections were collected using a diamond knife (Diatome Histo 6mm) and an ultramicrotome (Leica UC7). The blockface was carefully trimmed tight to the nerve in cross section using razor blades, and ultrathin sections (45-60nm) were transferred using a wire loop onto diced chips of silicon wafer, and dried on a hotplate set to 60°C. The silicon chips with adhered ultrathin sections were labeled with a diamond-tipped scribe and mounted on aluminum stubs using sticky carbon tabs before loading into the scanning electron microscope (SEM; Zeiss Sigma VP).

Images were collected using a backscattered electron detector (Gatan) at a working distance of approximately 6mm, with accelerating voltage set to 3kV, a 30µm aperture, and the beam in high current mode. Atlas5 control software (FIBICS) was used to capture a low magnification view of each section, and 5-7 high-resolution fields (4nm/px) of at least 20µm per side sampled from across the nerve cross section which were used for analysis.

### 3DEM imaging and visualization

Imaging of serial sections in the SEM (S3EM) was conducted as previously described^70, 71^, with some modifications. Briefly, samples used in 2D analysis from control and CalEx conditions were selected and blockfaces were trimmed using a 90° diamond trimming knife (Diatome) to a trapezoidal frustum of roughly 150×400 µm. A silicon chip (35×7mm; University Wafer, Boston, MA) was hydrophilized in a plasma cleaner (Harrick), rinsed in pure water, and partially immersed in a Diatome Histo knife, with one end sticking out of the water at the back of the boat. Four drops of pure ethanol were added to the water in the boat to attenuate surface tension, and an ionizing gun (Leica EM Crion) was activated and oriented towards the cutting edge of the knife mounted on the ultramicrotome. Ribbons of approximately 100 serial sections of a nominal 55 nm thickness were cut. When ribbons of sufficient quality and length were generated, they were released from the knife edge using a single-eyelash brush and carefully positioned over the chip. The water level was then slowly lowered, and sections were allowed to dry down on the silicon substrate over a few minutes. Chips were further dried on a hot plate set to 60°C for approximately 5 minutes and immediately labeled with a diamond scribe to indicate animal ID and nominal section thickness.

Samples were loaded into the same SEM, and imaged with the same imaging conditions using the array tomography software module of Atlas5. Briefly, low resolution (100nm/px) image maps of the ribbon(s) of serial sections were generated, and a mid-resolution (50nm/px) map of a central section was collected for evaluation. A region of interest (ROI) unobstructed by artifact throughout the series was identified and high resolution (8nm/px) images were collected from the ROI identified on consecutive sections. Following image intensity normalization in MIB2^72^, images were rigidly aligned using TrakEM2 in Fiji^73^, and fine alignment was accomplished using SWiFT-IR with compute resources provided from TACC through the 3dem.org portal^74^. The datasets for each animal constituted volumes of at least 15×15×4µm in dimension (with voxel sizes of 8×8×55nm).

Segmentation of myelin outfoldings and their corresponding axons was performed manually using the Volume Annotation and Segmentation Tool (VAST). The serial 2D segments were exported as .obj mesh files using the VAST Tools Matlab package. Meshes and data were imported into Blender software (blender.org) with the Neuromorph addon for further visualization^75^.

### TEM sample preparation

Transmission electron microscopy was completed in the Stanford Cell Sciences Imaging Facility. Samples were prepared according to previously published protocols (mobius protocol). Samples were washed in cold Karlsson-Schultz fixative (2.5% glutaraldehyde, 4% PFA in phosphate buffer, pH 7.3) and incubated in 2% OsO4 for four hours at 4°C with gentle shaking. The samples were then serially dehydrated in ethanol at 4°C and embedded in EmBed812 (EMS, 14120). 80 nm sections were taken using an UC7 (Leica, Wetzlar, Germany) and were collected on formvar/Carbon coated 100 mesh Cu grids. Sections were stained for 40 seconds in 3.5% uranyl acetate in 50% acetone followed by staining in Sato’s lead citrate for 2 minutes. Sections imaged in the JEOL JEM-1400 120kV. Images were taken using a Gatan OneView 4k X 4k digital camera. Quantification of TEM images was performed manually on Image-J.

### Statistics and analysis

Analysis of data in this paper was conducted blind to genotype and experimental condition (i.e. construct ID). Microsoft Excel 16.62.1 and GraphPad Prism 9.0 software were used for data analysis and statistical testing. Descriptive statistics (mean, SEM and biological replicates) and statistical testing were reported in Figure legends. To determine the method of statistical testing, q-q plots of each dataset were made to visualize whether the datasets were normally or non-normally distributed. For experiments performed with primary oligodendrocytes, statistics were performed on the means from biological replicates (cells from different cell preps); each mean was calculated from the technical replicates within a biological replicate. This distribution of data is represented in the superplots throughout the paper; each grey dot represents a value from a single cell, while colored dots represent means from biological replicates.

### Data and reagent availability

All constructs generated for this paper will be publicly available on AddGene. All raw data used in the manuscript will be deposited at Dryad at the time of acceptance for permanent storage and free access. All correspondence and requests for materials should be addressed to J.B.Z.

## Notes

### Competing Interest Statement

The authors have declared no competing interest.

### Summary of Updates

Minor update to title.

