## Supplemental Materials for "Oligodendrocyte calcium signaling sculpts myelin sheath morphology"

Contents include the following:

- Supplemental Figures and Figure Legends for S1-S7
- Legends for Supplemental Videos 1-3

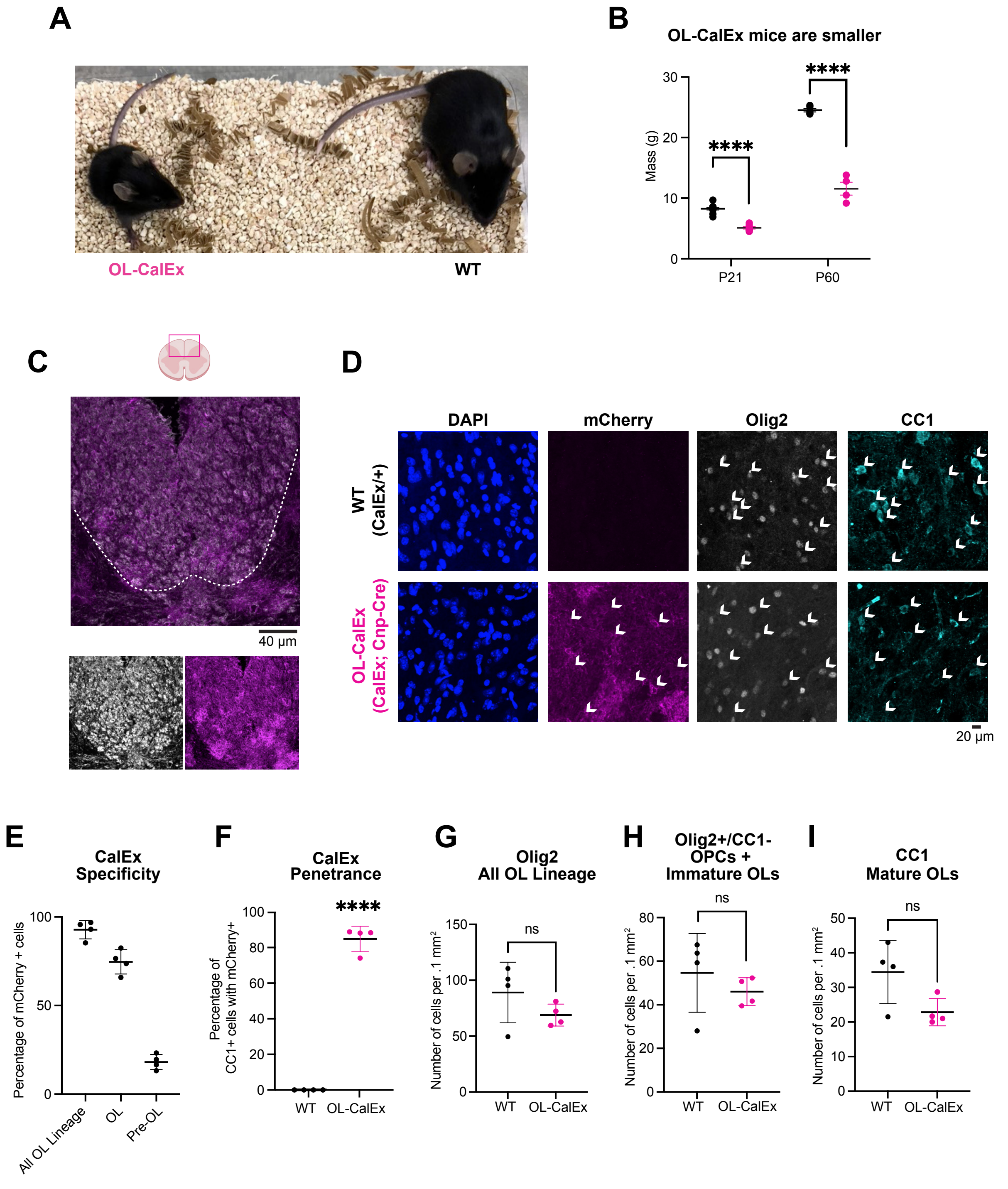


**Figure S1. CalEx is expressed in OL lineage cells and does not have overt effects on OL lineage numbers, related to Figure 1.**

(**A**) WT (right) and OL-CalEx (left) littermates at P60.
(**B**) Body weight for WT and OL littermates at P21 and P60. Statistical measurements (p-values) were determined by unpaired, two-tailed t-tests; ****$p < 0.0001$.
(**C**) Immunolabeling of P21 spinal cord cross section from OL-CalEx co-stained with MBP (grey) and mCherry (magenta). Dotted line denotes the border between white (dense, MBP rich regions on the edges of white matter) and grey matter. Scale bar, 40 μm. Created with Biorender.
(**D**) Confocal representative micrograph of control (top row) and OL-CalEx (bottom row) P21 dorsal spinal cords stained with DAPI (blue), mCherry (magenta), Olig2 (grey), and CC1 (teal). White arrows point out mature oligodendrocytes (Olig2+, CC1+ cells) Scale bar, 20 μm.
(**E**) Quantification of CalEx specificity by counting the percentage of mCherry+ cells with the following cell markers (average +/- SEM): Olig2 (all oligodendrocyte lineage cells), Olig2+/CC1+ (mature oligodendrocytes), and Olig2+/CC1- (immature oligodendrocytes and OPCs) N = 4 biological replicates.
(**F**) Quantification of CalEx penetrance by counting the percentage of mature CC1+ cells contained mCherry (average +/- SEM) in WT and OL-CalEx littermates. N = 4 biological replicates. Statistical measurement (p-value) was determined by an unpaired, two-tailed t-test; ****$p < 0.0001$.
(**G**) Quantification of number of all oligodendrocyte lineage cells per field of view in WT and OL-CalEx littermates. Three fields of view per biological replicate, N = 4 biological replicates. Statistical measurement (p-value) was determined by an unpaired, two-tailed t-test; n.s. not significant.
(**H**) Quantification of number of immature oligodendrocytes and OPCs per field of view in WT and OL-CalEx littermates. Three fields of view per biological replicate, N = 4 biological replicates. Statistical measurement (p-value) was determined by an unpaired, two-tailed t-test; n.s. not significant.
(**I**) Quantification of number of mature oligodendrocytes per field of view in WT and OL-CalEx littermates. Three fields of view per biological replicate, N = 4 biological replicates. Statistical measurement (p-value) was determined by an unpaired, two-tailed t-test; n.s. not significant.

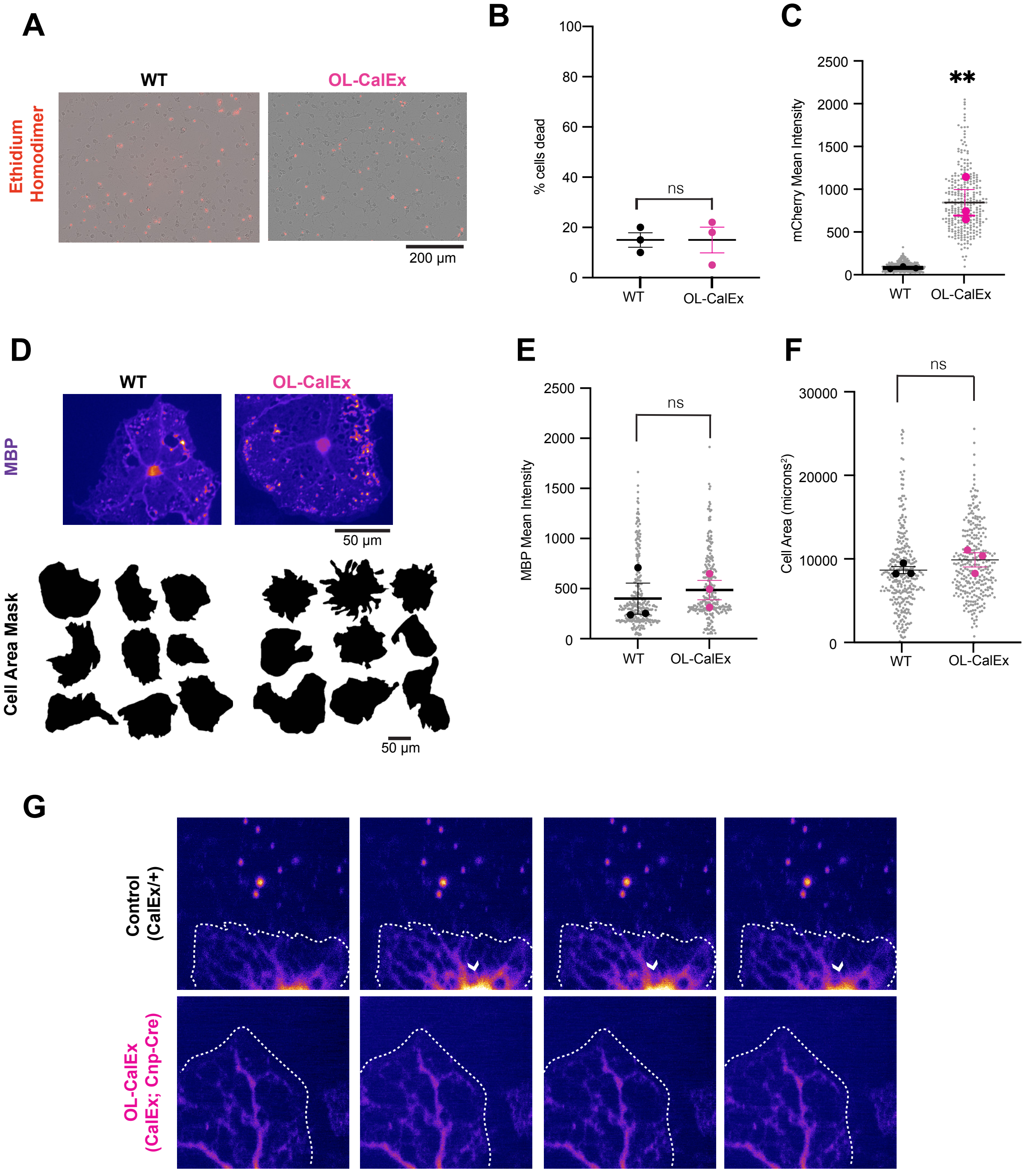


**Figure S2. Characterization of CalEx in primary cultured oligodendrocytes, related to Figure 1.**

(**A**) Representative micrograph of WT and OL-CalEx treated cells. Cells visualized using brightfield and ethidium homodimer is depicted in red overlay. Scale bar, 200 μm

(**B**) Quantification of percentage of dead cells (Average +/- SEM, 3 FOV per condition, N = 3 biological replicates/preps) in WT and OL-CalEx

(**C**) Quantification of mCherry fluorescence in WT versus OL-CalEx cells. N = 3 biological replicates. Statistical measurement (p-value) was determined by an unpaired, two-tailed t-test; **$p < 0.01$.

(**D**) (top) Representative micrograph of myelin basic protein staining in (top) WT and (bottom) OL-CalEx oligodendrocytes. (bottom) Representative cell area masks for WT or OL-CalEx cells. Scale bar, 50 μm.

(**E**) Quantification of myelin basic protein mean intensity (Average +/- SEM) in WT and OL-CalEx oligodendrocytes. Statistical measurement (p-value) was determined by an unpaired, two-tailed t-test; n.s. not significant.

(**F**) Quantification of cell area (Average +/- SEM) for WT or OL-CalEx cells. Statistical measurement (p-value) was determined by an unpaired, two-tailed t-test; n.s. not significant.

(**G**) Representative micrograph and traces of calcium transient in WT oligodendrocyte process (top) and lack of calcium fluctuations in OL-CalEx oligodendrocyte (bottom). White arrowhead points to calcium transient occurring in the soma of the cultured oligodendrocyte. White dotted lines denote cell borders.

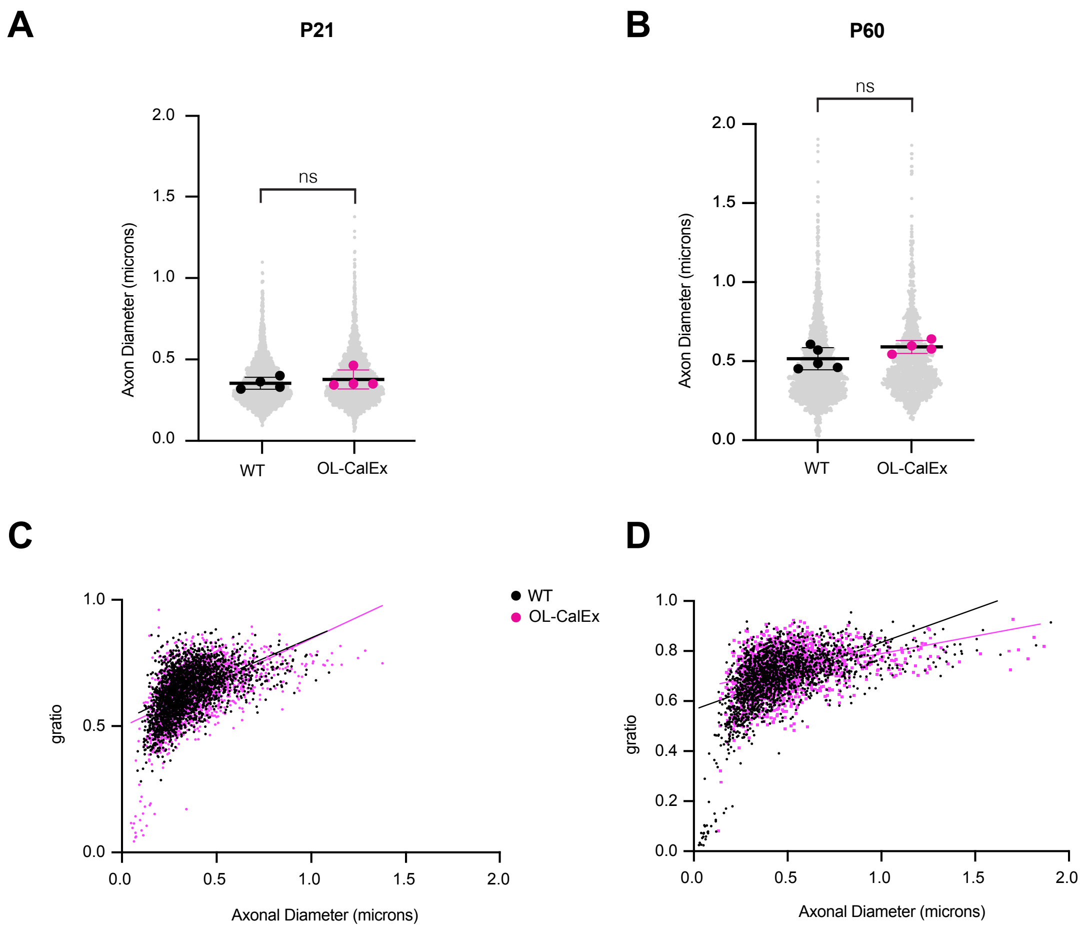


**Figure S3. Additional ultrastructural analysis of OL-CalEx and WT littermate mice, related to Figure 1.**

(**A**) Quantification of axonal diameter in P21 optic nerves. Average +/- SEM of WT and OL-CalEx littermates. N = 4 biological replicates Statistical measurement (p-value) was determined by an unpaired, two-tailed t-test; n.s. not significant.

(**B**) Quantification of axonal diameter in P60 optic nerves. Average +/- SEM of WT N = 5 and OL-CalEx littermates N = 4. Statistical measurement (p-value) was determined by an unpaired, two-tailed t-test; n.s. not significant.

(**C**) Distribution of *g*-ratio versus axonal caliber in WT vs OL-CalEx optic nerves at P21.

(**D**) Distribution of *g*-ratio versus axonal caliber in WT vs OL-CalEx optic nerves at P60.

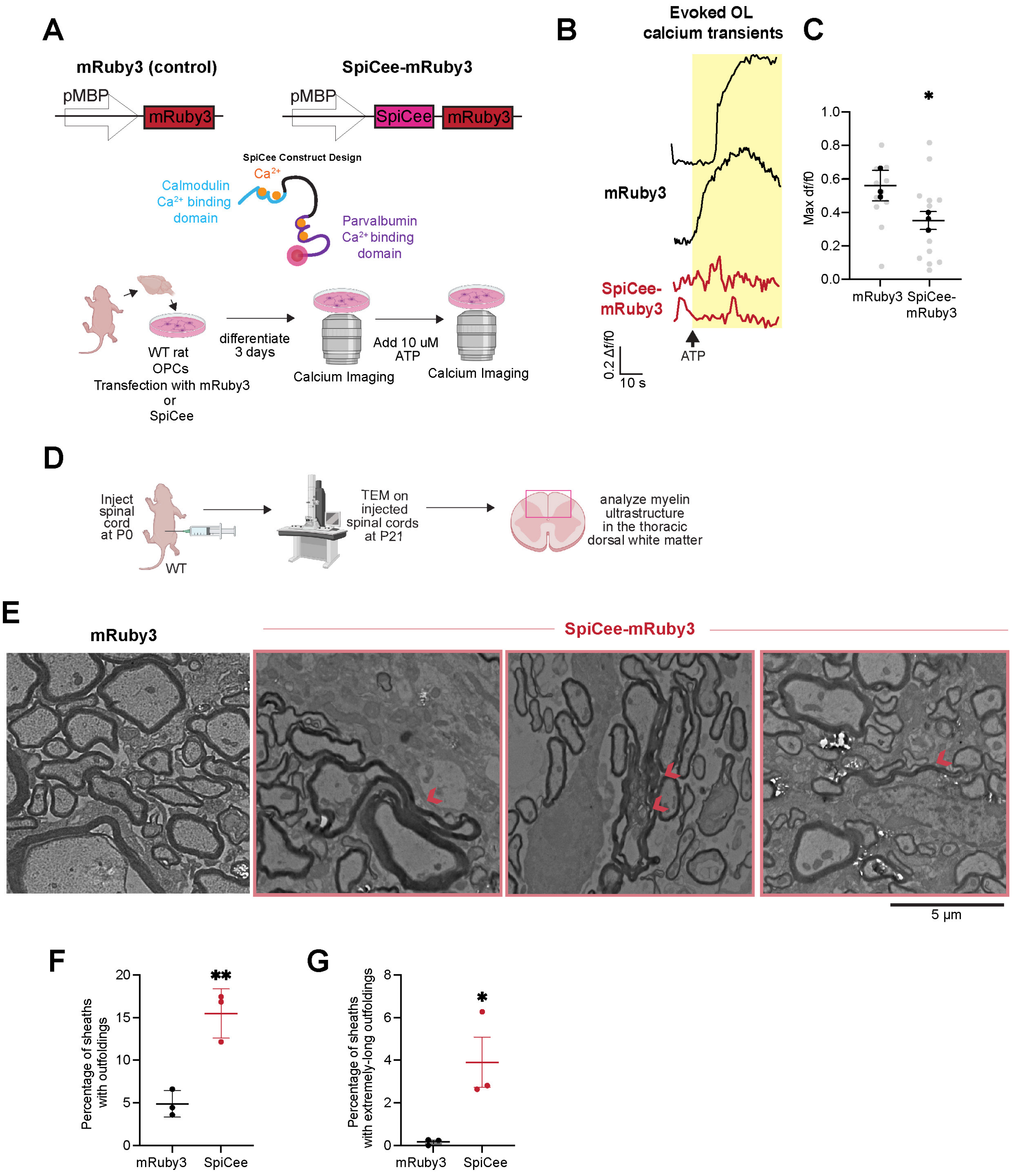


**Figure S4. Oligodendrocyte-specific expression of SpiCee "calcium sponge" induces outfolding formation in vivo, related to Figure 2.**

**(A)** Construct design for MBP promoter driven mRuby3 (control, left) and MBP promoter driven SpiCee mRuby3 (right). SpiCee is made up of a low-$Ca^{2+}$-affinity calmodulin domain and a high $Ca^{2+}$-affinity parvalbumin domain joined by a floppy linker and mRuby3 downstream to label cells that express SpiCee. SpiCee was validated by transfection of OPCs with either mRuby3 or SpiCee-mRuby3. Cells differentiated for 3 days before performing calcium imaging. To stimulate calcium transients in oligodendrocytes ATP was added to the cells.

**(B)** Example traces from (top) or SpiCee transfected oligodendrocytes. Arrow indicates addition of ATP.

**(C)** Quantification of maximum amplitude of calcium transients after ATP addition. Average +/- SEM, N = 3 technical replicates. P-value determined by unpaired, two-tailed Student’s t-test; *$p < 0.05$.

**(D)** Virus encoding for SpiCee-mRuby3 or mRuby3 was injected into P0 mouse pups. Injected spinal cords were harvested at P21 and processed for transmission electron microscopy. Created with Biorender.com

**(E)** Example TEM micrograph of mRuby3 injected (left) or SpiCee-mRuby3 injected (right) P21 spinal cords. Red arrows point to outfoldings. Scale bar, 5 μm.

**(F)** Quantification of percentages of myelin sheaths with outfoldings at P21 in e. Average +/- SEM, P21, N = 3. p-value determined by unpaired, two-tailed Student’s t-test; **$p < 0.01$.

**(G)** Quantification of percentages of myelin sheaths with extremely-long outfoldings (defined as an outfolding greater than 2x the diameter of the axons) in e. Average +/- SEM, P21, N = 3. p-value determined by unpaired, two-tailed Student’s t-test; *$p < 0.05$.

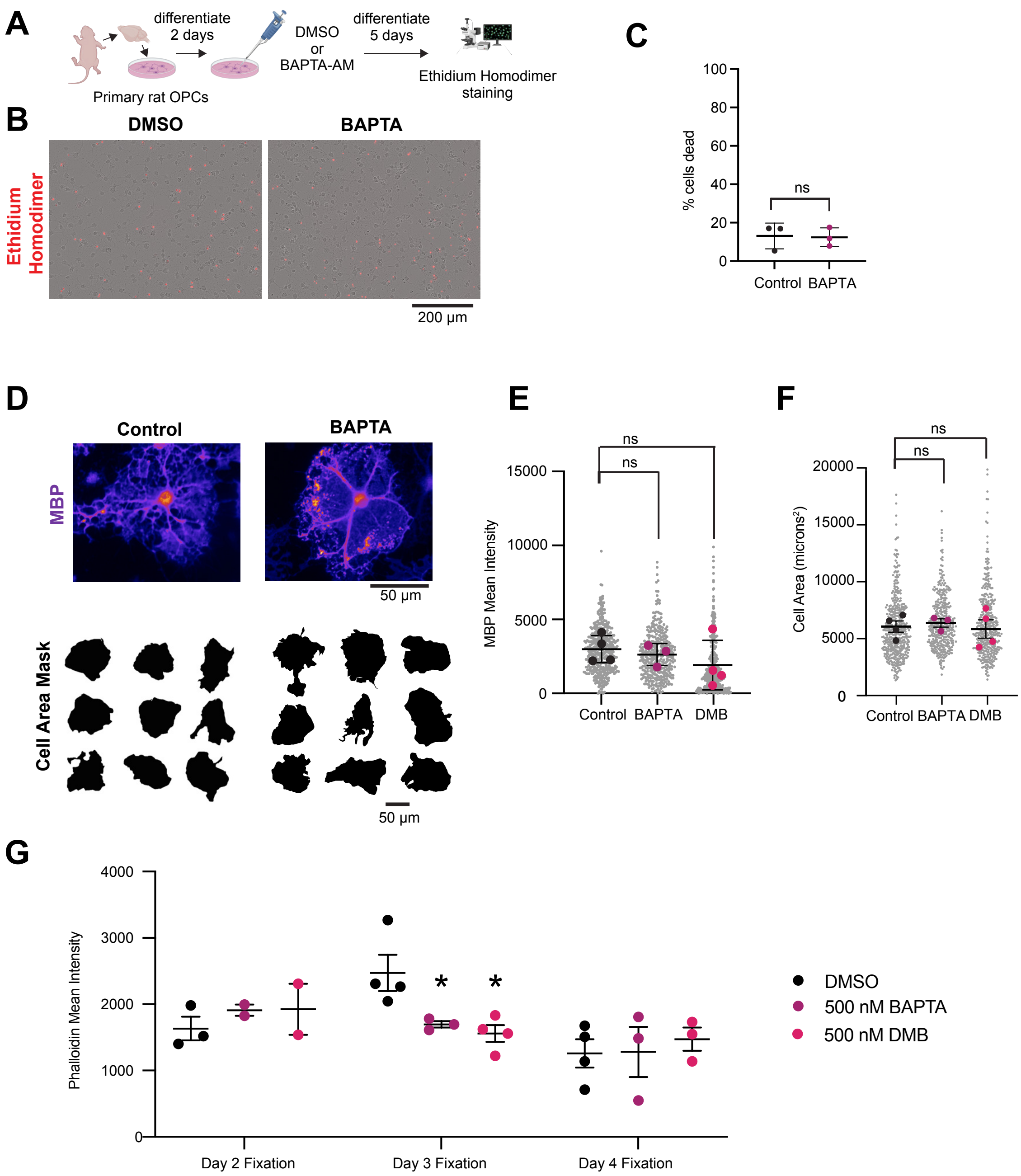


**Figure S5. Additional data on BAPTA/DMB treatment of primary cultured oligodendrocytes, related to Figure 3.**

(**A**) Experimental set up for determining percentage of dead cells in BAPTA treated cells. Primary rat OPCs were differentiated for two days and then treated overnight with either DMSO or BAPTA-AM. Cell media was then replaced with normal OL media and cells continued differentiating for 5 days before treatment with ethidium homodimer to label dead cells. Scale bar, 200 μm.

(**B**) Representative micrograph of DMSO and BAPTA treated cells. Cells visualized using brightfield and ethidium homodimer is depicted in red overlay.

(**C**) Quantification of percentage of dead cells (Average +/- SEM, 3 FOV per condition, N = 3 biological replicates/preps) in DMSO-treated and BAPTA-treated cells; P-value determined by unpaired, two-tailed Student's t-test; n.s., not significant.

(**D**) Representative micrograph of myelin basic protein staining (top) and (bottom) cell area masks for cells treated with DMSO or BAPTA. Scale bar, 50 μm.

(**E**) Quantification of myelin basic protein mean intensity (Average +/- SEM) in DMSO treated, BAPTA-AM treated, or DMB-AM treated. P-value determined by one way ANOVA; n.s., not significant.

(**F**) Quantification of cell area (Average +/- SEM) in DMSO treated, BAPTA-AM, and DMB-AM treated. P-value determined by unpaired, two-tailed Student's t-test; n.s., not significant.

(**G**) Quantification of phalloidin mean intensity (Average +/- SEM) across three days of oligodendrocyte differentiation; P-value determined by one-way ANOVA;*$p < 0.05$.

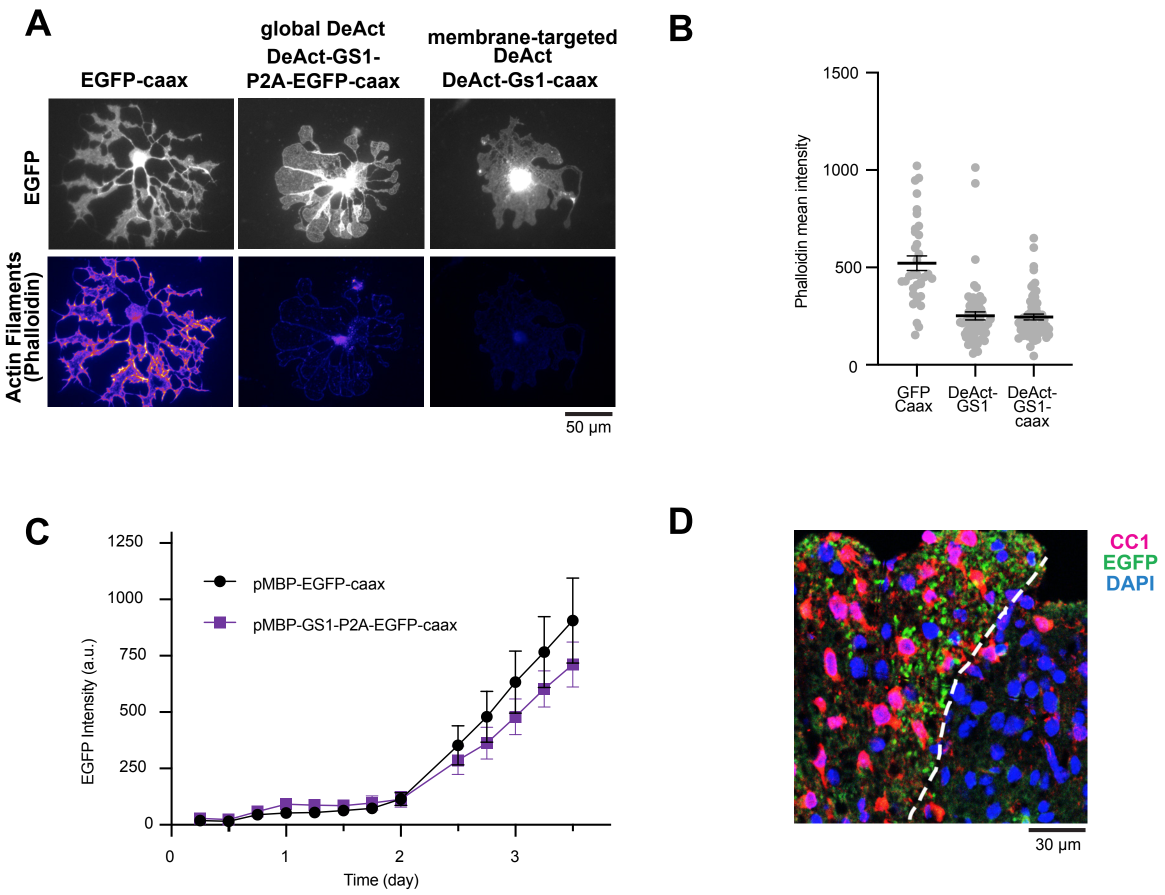


**Figure S6. Validation of DeActs in cultured and in vivo oligodendrocytes, related to Figure 4.**

(**A**) Representative micrograph of phalloidin staining (bottom) and EGFP expression (top) for cells transfected with either EGFP-caax, DeAct-GS1, or DeAct-GS1-caax. Scale bar, 50 μm.

(**B**) Quantification of phalloidin mean intensity (Average +/- SEM) in EGFP-caax, DeAct-GS1 or DeAct-GS1-caax.

(**C**) Time course of expression of MBP promoter (pMBP)-driven expression of EGFP-caax and GS1-P2A-EGFP-caax.

**(D)** Confocal micrograph of spinal cord cross section injected with EGFP-caax. CC1: Magenta, EGFP: Green, DAPI: Blue. Dotted line denotes the border of the white matter and grey matter. Scale bar, 30 μm

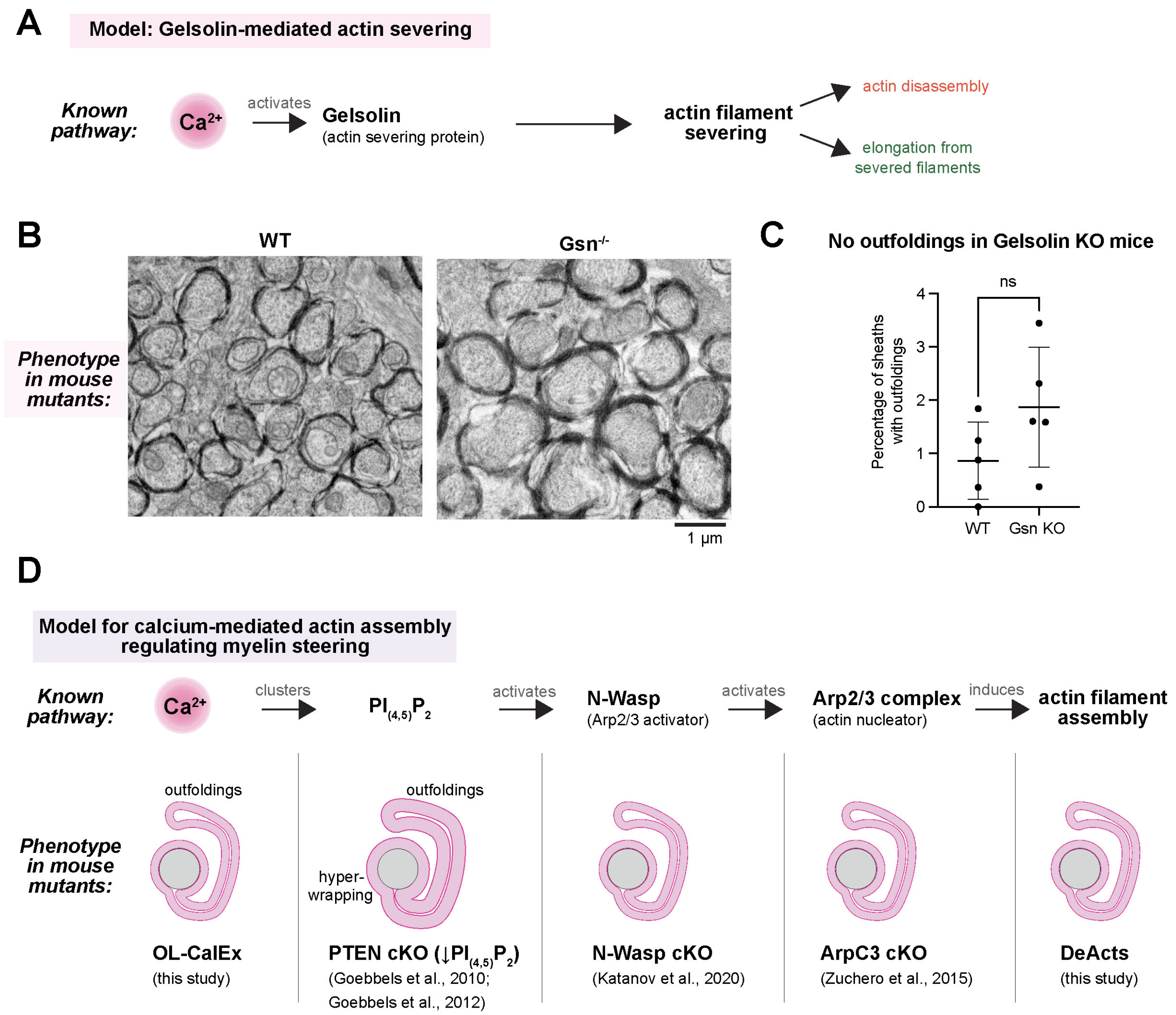


**Figure S7. Alternative model for how calcium may regulate actin severing in developing myelin sheaths, related to Figure 6.**

(**A**) Model for how calcium may regulate actin severing in oligodendrocytes to steer myelin membrane growth. Calcium is known to activate actin-severing protein gelsolin, which promotes both actin disassembly as well as formation of new filaments from newly-generated barbed ends.

(**B**) Transmission Electron Microscopy WT (left) and Gsn KO (right) optic nerve sections. Scale bar, 1 µm.

(**C**) Quantification of myelin sheaths with outfoldings in e. Average +/- SEM, N = 5. P-value determined by unpaired, two-tailed t-test; n.s., not significant.

(**D**) Molecular mechanism for how calcium may regulate actin assembly. Calcium is known to cluster PIP2, which in turn activates N-WASP to activate the Arp2/3 complex to induce actin filament assembly. Conditional knockout of any component of this pathway leads to increased outfoldings.

### SUPPLEMENTAL VIDEOS

**Supplemental Video 1. Calcium signaling in WT and OL-CalEx primary oligodendrocytes, related to Figure 1.** (right) Fluo4-AM loaded primary WT oligodendrocyte at day 3 of differentiation. (left) Fluo4-AM loaded primary OL-CalEx oligodendrocyte at day 3 of differentiation.

**Supplemental Video 2. 3-dimensional reconstruction of two myelinated axons with outfoldings from OL-CalEx optic nerve, related to Figure 2.**

**Supplemental Video 3. 360-degree view of 3-dimensional reconstruction of two myelin sheaths with outfoldings from OL-CalEx optic nerve, related to Figure 2.**
